# The evolutionary history of ACE2 usage within the coronavirus subgenus *Sarbecovirus*

**DOI:** 10.1101/2020.07.07.190546

**Authors:** H.L Wells, M Letko, G Lasso, B Ssebide, J Nziza, D.K Byarugaba, I Navarrete-Macias, E Liang, M Cranfield, B.A Han, M.W Tingley, M Diuk-Wasser, T Goldstein, C.K Johnson, J Mazet, K Chandran, V.J Munster, K Gilardi, S.J Anthony

## Abstract

SARS-CoV-1 and SARS-CoV-2 are not phylogenetically closely related; however, both use the ACE2 receptor in humans for cell entry. This is not a universal sarbecovirus trait; for example, many known sarbecoviruses related to SARS-CoV-1 have two deletions in the receptor binding domain of the spike protein that render them incapable of using human ACE2. Here, we report three sequences of a novel sarbecovirus from Rwanda and Uganda which are phylogenetically intermediate to SARS-CoV-1 and SARS-CoV-2 and demonstrate via in vitro studies that they are also unable to utilize human ACE2. Furthermore, we show that the observed pattern of ACE2 usage among sarbecoviruses is best explained by recombination not of SARS-CoV-2, but of SARS-CoV-1 and its relatives. We show that the lineage that includes SARS-CoV-2 is most likely the ancestral ACE2-using lineage, and that recombination with at least one virus from this group conferred ACE2 usage to the lineage including SARS-CoV-1 at some time in the past. We argue that alternative scenarios such as convergent evolution are much less parsimonious; we show that biogeography and patterns of host tropism support the plausibility of a recombination scenario; and we propose a competitive release hypothesis to explain how this recombination event could have occurred and why it is evolutionarily advantageous. The findings provide important insights into the natural history of ACE2 usage for both SARS-CoV-1 and SARS-CoV-2, and a greater understanding of the evolutionary mechanisms that shape zoonotic potential of coronaviruses. This study also underscores the need for increased surveillance for sarbecoviruses in southwestern China, where most ACE2-using viruses have been found to date, as well as other regions such as Africa, where these viruses have only recently been discovered.

## Introduction

The recent emergence of *severe acute respiratory syndrome coronavirus 2* (SARS-CoV-2) in China and its rapid spread around the world demonstrates that coronaviruses (CoVs) from wildlife remain an urgent threat to global public health and economic stability. In particular, coronaviruses from the subgenus *Sarbecovirus* (which includes SARS-CoV-2, SARS-CoV-1, numerous bat viruses, and a small number of pangolin viruses) [1] are considered to be a high-risk group for potential emergence. As both sarbecoviruses that have caused human disease (SARS-CoV-1 and −2) use angiotensin-converting enzyme 2 (ACE2) as their cellular receptor [2,3], the evolution of this trait is of particular importance for understanding the emergence pathway for sarbecoviruses. Bat SARS-like coronavirus Rp3 is a phylogenetically close relative of SARS-CoV-1 but is unable to bind human ACE2 (hACE2) *in vitro* [4]. In contrast, other close relatives of SARS-CoV-1, including bat SARS-like coronavirus WIV1 and WIV16, do have the capacity to bind hACE2 [5,6]. A number of other SARS-CoV-1-like viruses have also been tested for their ability to utilize hACE2 [7–9] and comparison of their spike protein sequences shows that viruses that are unable to utilize hACE2 unanimously have one or two deletions in their RBDs that make them structurally very different than those that do use hACE2 [8]. As SARS-CoV-1, Rp3, WIV1, and WIV16 viruses are closely phylogenetically related, the evolutionary mechanism explaining the variation in their ability to utilize hACE2 (and likely also bat ACE2) as a cellular receptor has thus far been unclear.

Chinese horseshoe bats (*Rhinolophidae*) are thought to be the primary natural reservoir of sarbecoviruses [5,7,10–12]. Bats within this family are also considered to be the source of the progenitor virus to SARS-CoV-1, as related viruses with high sequence identity to SARS-CoV-1 have been sequenced from Rhinolophid bats, although none have high sequence similarity to SARS-CoV-1 across the entire genome [7,13]. It is hypothesized that SARS-CoV-1 obtained genomic regions from different strains of bat SARS-1-like CoVs in or near Yunnan Province by recombination before spilling over into humans [7,13,14]. In particular, one region of SARS-CoV-1 that is known to have a recombinant origin is the spike gene, as a breakpoint has been detected at the junction of ORF1b and the spike [13,15]. The SARS-1-CoV spike is genomically very different from other viruses in the same clade that have large deletions in the receptor binding domain (RBD) and are unable to use hACE2. The exact minor parent that contributed the recombinant region is still unknown, but it was previously hypothesized that the recombination occurred with a yet undiscovered lineage of sarbecoviruses and that this event contributed strongly to its potential for emergence [13,16]. Recombination has also been shown within the spike genes of other CoVs that have spilled over into humans and domestic animals and is potentially an important driver of emergence for all coronaviruses [17–22].

In order for CoVs to recombine, they must first have the opportunity to do so by sharing overlapping geographic ranges, host species tropism, and cell and tissue tropism. Sarbecoviruses in bats tend to phylogenetically cluster according to the geographic region in which they were found [7,23]. Yu et. al showed that there are three lineages of SARS-CoV-1-like viruses: Lineage 1 from southwestern China (Yunnan, Guizhou, and Guangxi, and including SARS-CoV-1), Lineage 2 from other southern regions (Guangdong, Hubei, Hong Kong, and Zhejiang), and Lineage 3 from central and northern regions (Hubei, Henan, Shanxi, Shaanxi, Hebei, and Jilin) [23]. Studies in Europe and Africa have shown that there are distinct sarbecovirus clades in each of these regions as well, herein named “Lineage 4” [24–29]. Sarbecoviruses appear to switch easily among co-occurring *Rhinolophus* species [30,31]; however, they appear to rarely occupy more than one geographic area, despite the fact that some of these bat species have widespread distributions across China.

Shortly after the emergence of SARS-CoV-2, Zhou et al. showed a high degree of homology across the genome between a bat virus (RaTG13) sampled from Yunnan Province in 2013 and SARS-CoV-2 [3]. RaTG13 has also been shown to bind hACE2, although with decreased affinity compared to SARS-CoV-2 [32]. Subsequently, seven full- or near full-length SARS-CoV-2-like viruses were published that had been sampled from Malayan pangolins (*Manis javanica*) in 2017 and 2019 [33,34], one of which has also been tested and found to bind hACE2 [35]. Neither SARS-CoV-2, RaTG13, nor the pangolin CoVs have deletions in their RBDs. In contrast, the most recently described bat virus (RmYN02) is even more closely related to SARS-CoV-2 than RaTG13 in the polymerase gene and was also found in Yunnan Province; however, this sequence has deletions in the RBD and homology modeling suggests it likely does not use hACE2 [36]. Together, these viruses form a fifth phylogenetic lineage (“Lineage 5”) that is distinct from all other lineages of sarbecoviruses despite having been detected in Yunnan, where all viruses found until this point had belonged to Lineage 1.

This finding of overlapping Lineage 1 and Lineage 5 viruses in geographic space is inconsistent with the previously observed pattern of biogeography for sarbecoviruses. SARS-CoV-2 was isolated first from people in Hubei Province and one of the pangolin viruses was isolated from an animal sampled in Guangdong, neither of which are Lineage 1 provinces. However, the true geographic origins of these viruses are unknown as it is possible they were anthropogenically transported to the regions in which they were detected. For example, the Malayan pangolin (*Manis javanica*) has a natural range that reaches southwestern China (Yunnan Province) at its northernmost edge and extends further south into Myanmar, Lao PDR, Thailand, and Vietnam [37]. So, if they were naturally infected (as opposed to infection via wildlife trade), the infection was potentially not acquired from Guangdong Province. Similarly, SARS-CoV-2 cannot be guaranteed to have emerged from bats in Hubei Province, as humans are highly mobile and the exact spillover event was not observed. If the clade containing SARS-CoV-2 and its close relatives is indeed endemic in animals in Yunnan and the nearby Southeast Asian regions as suggested by the presence of RaTG13, RmYN02, and the natural range of the Malayan pangolin, whatever mechanism is facilitating the biogeographical concordance of Lineages 1, 2 and 3 within China appears to no longer apply for the biogeography of Lineage 5, since they all appear to overlap in and around Yunnan Province.

Here, we report a series of observations that together suggest that SARS-CoV-1 and its close relatives gained the ability to utilize ACE2 through a recombination event that happened between an ancestor of SARS-CoV-1 and a Lineage 5 virus phylogenetically related to SARS-CoV-2, which could only have occurred with the lineages occupying the same geographic and host space. We also report three full-length genomes of sarbecoviruses from Rwanda and Uganda and demonstrate that the RBDs of these viruses are genetically intermediate between viruses that use ACE2 and those that do not. Accordingly, we also investigate the potential for these viruses to utilize hACE2 *in vitro*. Together, our findings help illuminate the evolutionary history of ACE2 usage within sarbecoviruses and provide insight into identifying their risk of emergence in the future. We also propose a mechanism that could explain the pattern of phylogeography across Lineages 1, 2, and 3, and why Lineage 5 viruses (including SARS-CoV-2 and its relatives) represent an inconsistency to this pattern.

## Results

To better understand the evolutionary history of sarbecoviruses we first constructed a phylogenetic tree of the RNA-dependent RNA polymerase (RdRp) gene, also known as nsp12 (Figure 1). The tree was constructed using sequences from GenBank as well as three sequences of a novel sarbecovirus detected in bats from Uganda and Rwanda as part of the USAID-PREDICT project. The three novel sequences share >99% nucleotide identity to each other and ~76% and ~74% nucleotide identity with SARS-CoV-1 and SARS-CoV-2, respectively. Phylogenetically, they lie within Lineage 4, clustering with previously reported SARS-related coronavirus BtKY72 found in bats in Kenya [29] and bat coronavirus BM48-31 from Bulgaria [26]. The topology of the sarbecovirus phylogeny is uncertain with respect to the placement of the Lineage 4 viruses, with some models placing them between Lineage 5 and Lineages 1, 2, and 3, and others placing them at the base of the tree, depending on the methodology and alignment used [3,38,39] (Supplementary Figure S1). Our results place Lineage 4 in the former position with high posterior support for the RdRp gene, though the variability in this placement must be recognized. Figure 1 also demonstrates the same geographic pattern of concordance reported by Yu et al [23], where viruses in each lineage show a clear pattern of fidelity with particular geographic regions. However, SARS-CoV-2 does not lie within the clade of bat sarbecoviruses that have been detected in bats in China to date but rather forms a much deeper, separate lineage. The discovery of the “Lineage 5” clade containing SARS-CoV-2 and related viruses in pangolins and bats is a deviation from the geographic patterns observed for other sarbecoviruses.

**Figure 1:**
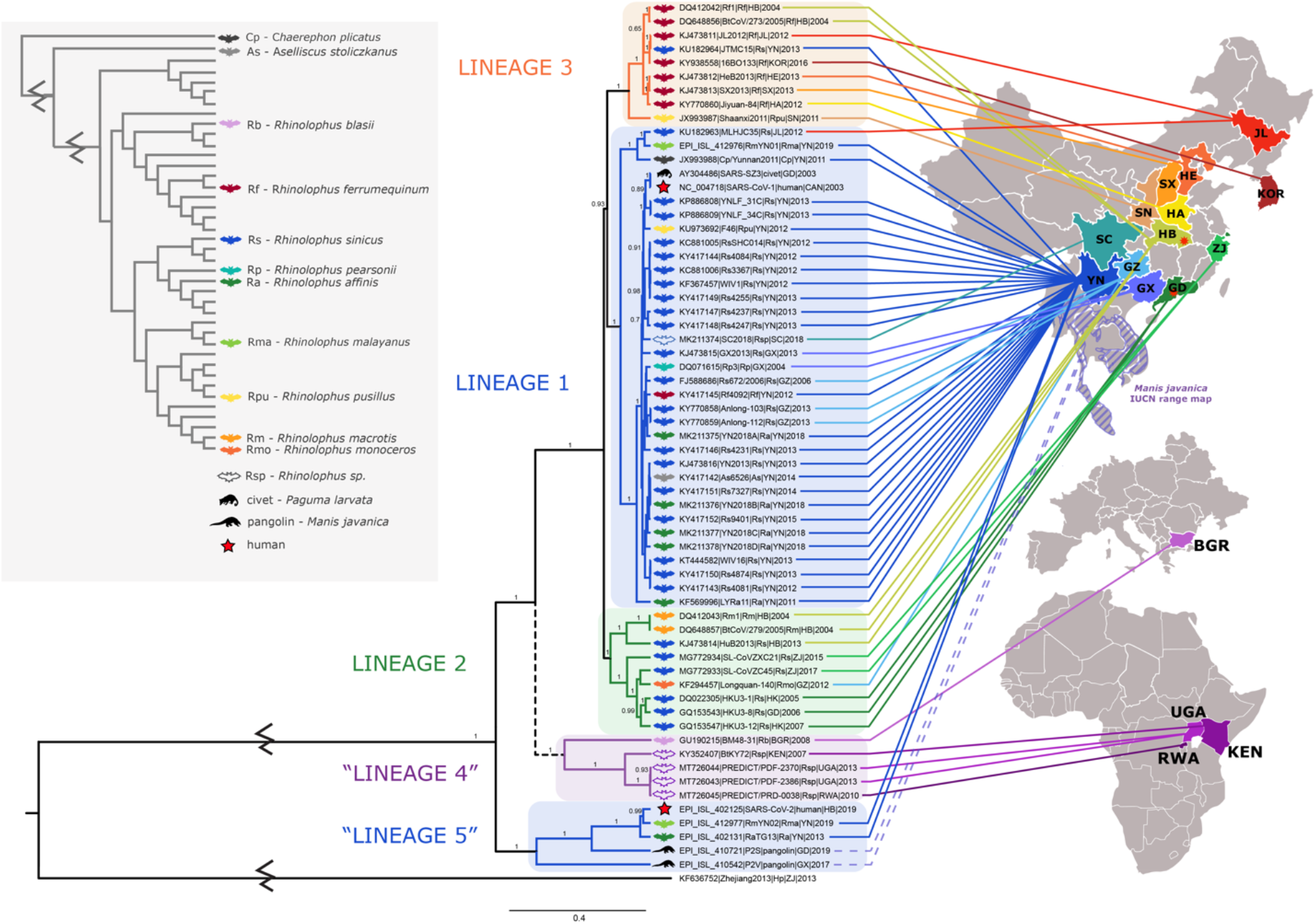
Phylogenetic tree of the RNA dependent RNA polymerase (RdRp) gene (nsp12) and associated geographic origin and host species. Colors of clade bars represent the different geographic lineages. Lineage 1 is shown in blue, Lineage 2 in green, and Lineage 3 in orange. The clade of viruses from Africa and Europe is putatively named “Lineage 4” and is shown in purple. The phylogeny shows strong posterior support for the branching order presented; however, different models or genes have produced trees with different branching orders placing Lineage 4 outside Lineage 5, so the branch to Lineage 4 is dashed to represent this uncertainty (Supplementary Figure S1). The putative “Lineage 5” containing SARS-CoV-2 is also shown in blue at the bottom of the tree to demonstrate that the sequences are from the same regions as Lineage 1 viruses. The geographic origin of each virus is indicated by the lines that terminate in the respective country or province with the same color code. The full province and country names for all two- and three-letter codes can be found in Table 1. As human, civet, and pangolin viruses cannot be certain to have naturally originated in the province in which they were first found, their locations are not illustrated, but the natural range of the pangolin (*Manis javanica*) is denoted with dashed shading and the origins of the SARS-CoV-1 and SARS-CoV-2 human outbreaks are designated with red stars in Guangdong and Hubei, respectively. Hosts are also shown with colored symbols according to the key on the left. The host phylogeny in the key was adapted from [81]. The root of the tree was shortened for clarity.

To investigate the evolutionary history of ACE2 usage, we built a second phylogenetic tree using only the RBD of the spike gene and compared it to the phylogeny of RdRp (Figure 2). This region was selected because the spike protein mediates cell entry and because previous reports showed that SARS-CoV-1 and SARS-CoV-2 both use hACE2, despite being distantly related in the RdRp [2,3]. Within the RBD region of the genome, SARS-CoV-1 and all ACE2-using viruses are much more closely related to SARS-CoV-2 than to other Lineage 1 viruses (Figure 2). Interestingly, bat virus RmYN02 is no longer associated with SARS-CoV-2 in the RBD and is instead within the clade of non-ACE2-using viruses. We also found that within the RBD, ACE2-using viruses and non-ACE2-using viruses are perfectly phylogenetically separated. The viruses from Africa and Europe form a distinct clade that is intermediate between the ACE2-using and non-ACE2-using groups, but appears more closely related to the ACE2-using group.

**Figure 2:**
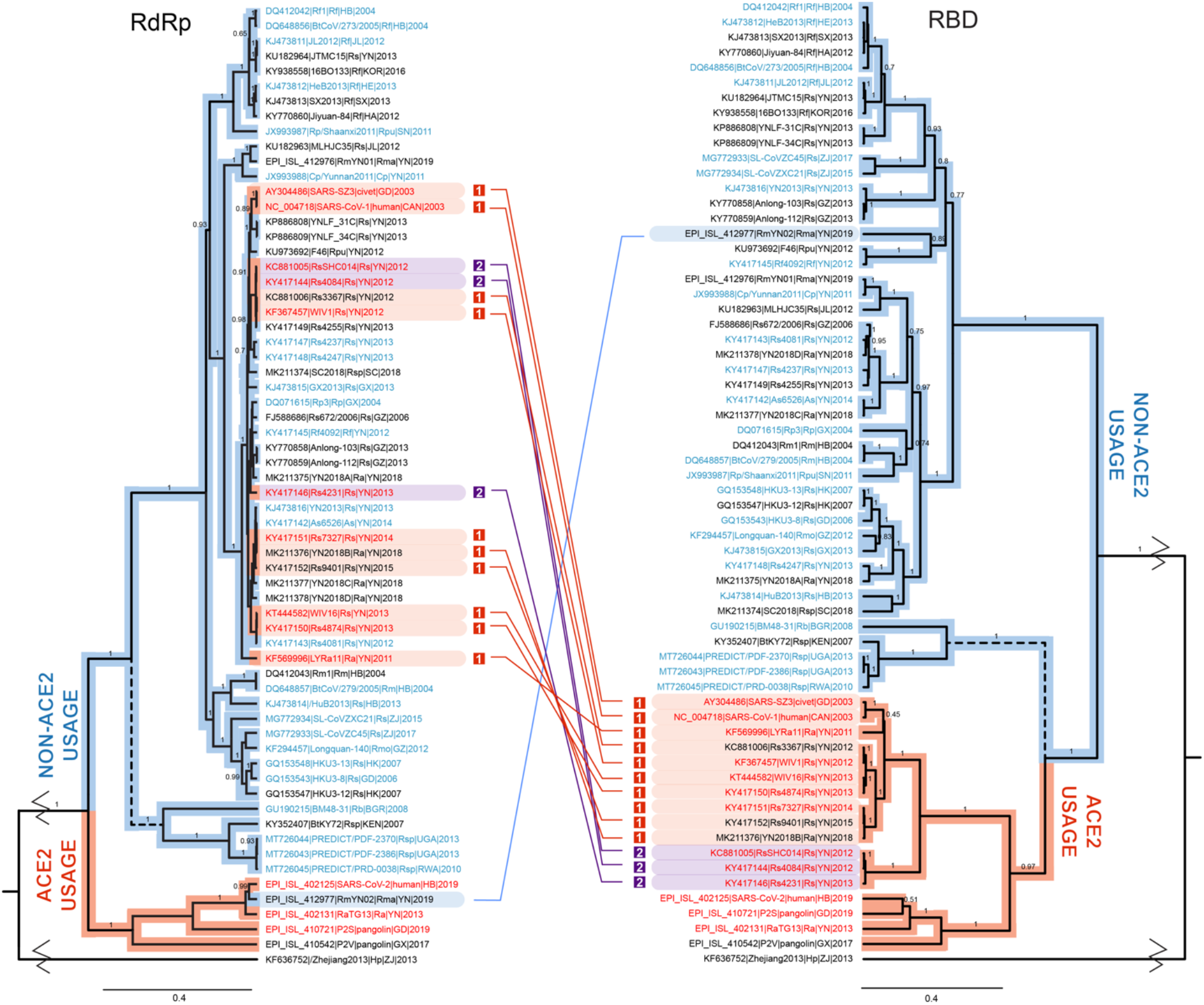
Phylogenetic trees of RdRp (left) and the RBD (right) demonstrating recombination events between ACE2-users and non-ACE2-users. Names of viruses that have been confirmed to use hACE2 are shown in red font, and those that have been shown to not use hACE2 are shown in blue font (citations can be found in Table 1). Viruses in black font have not yet been tested. The red and blue highlighted clade bars separate viruses with the structure associated with ACE2 usage (highly similar to viruses confirmed to use hACE2 specifically) and the structure with deletions that cannot use ACE2, respectively. Connecting lines indicate recombination events that resulted in a gain of ACE2 usage (red) or a loss of ACE2 usage (blue). The two different groups of RBD sequence within the Lineage 1 recombinants that gained ACE2 usage are distinguished in red (Type 1) and purple (Type 2) highlighting. The distances of the roots have been shortened for clarity. The branch leading to Lineage 4 is dashed to demonstrate uncertainty in its positioning.

While these viruses from Africa and Europe are slightly more similar to the ACE2-using group, they differ somewhat in amino acid sequence from the ACE2-users at the binding interface, including a small deletion in the middle of the sequence (Figure 5, region 2). Thus, to determine the ability of these sarbecoviruses to use hACE2 and better delineate the boundaries of ACE2 usage, we performed *in vitro* experiments in which we replaced the RBD of SARS-CoV-1 with the RBD from the Uganda (PDF-2370, PDF-2386) and Rwanda viruses (PRD-0038) [8]. Single-cycle Vesicular Stomatitis Virus (VSV) reporter particles containing the recombinant SARS-Uganda and SARS-Rwanda spike proteins were then used to infect BHK cells expressing hACE2. While VSV-SARS-CoV-1 showed efficient usage of hACE2, VSV-Uganda and VSV-Rwanda did not (Figure 3).

**Figure 3:**
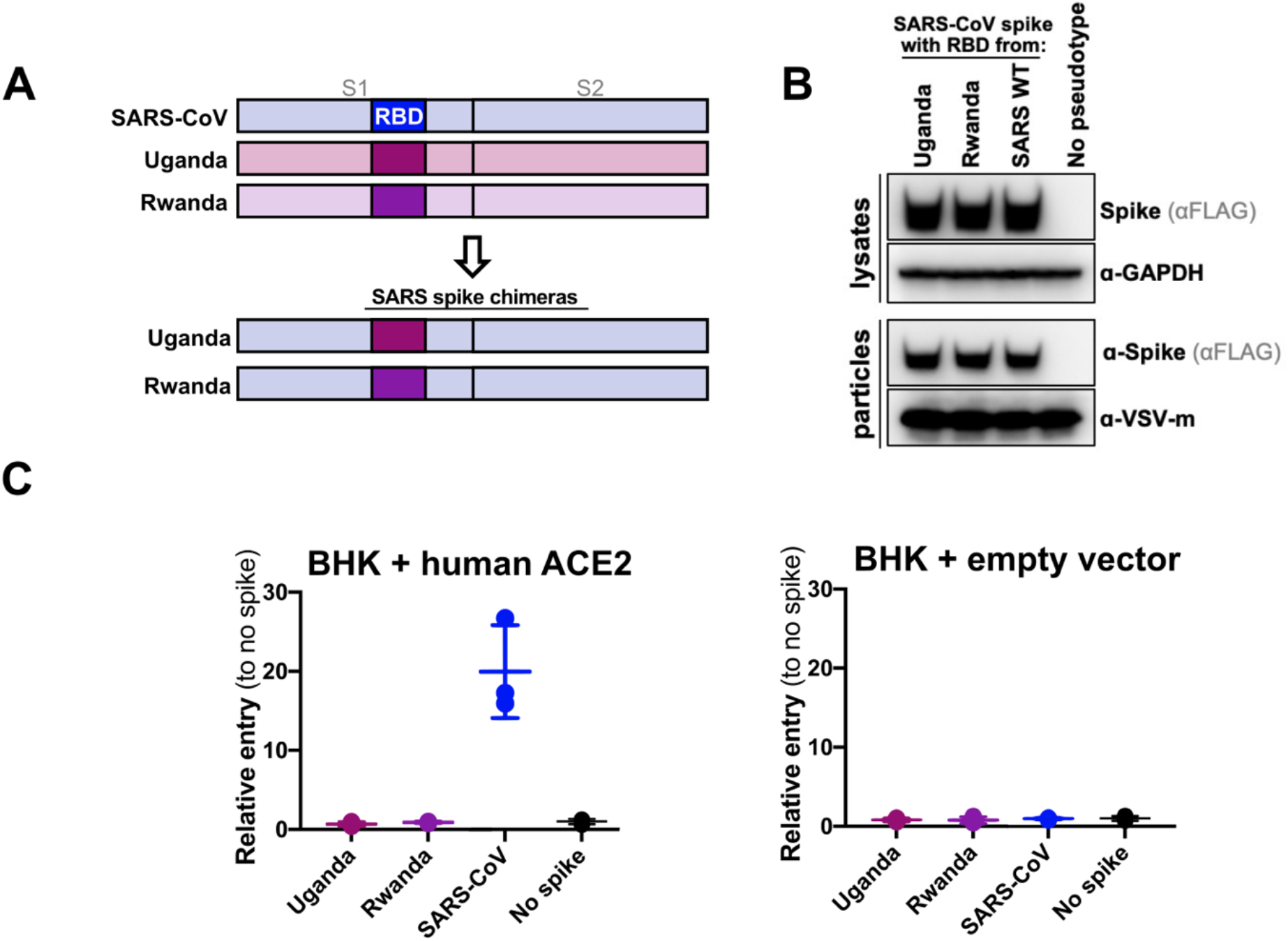
hACE2 usage of bat sarbecoviruses investigated using a surrogate VSV-psuedotyping system. (A) Schematic showing the structure of chimeric spike proteins. The SARS-CoV-1 spike backbone is used in conjunction with the RBD from the Uganda and Rwanda strains. (B) Incorporation of chimeric SARS-CoV-1 spike proteins into VSV. Western blots show successful expression of chimeric spikes (lysates) and their incorporation into VSV (particles). (C) hACE2 entry assays. Left, wildtype SARS-CoV spike protein is able to mediate entry into BHK cells expressing hACE2. In contrast, recombinant spike proteins containing either the Uganda or Rwanda RBD were unable to mediate entry. Entry is expressed relative to VSV particles with no spike protein. Right, control experiment for entry assay. BHK cells do not express hACE2 and therefore do not permit entry of hACE2-dependent VSV pseudotypes.

To try and explain why the African sarbecoviruses are unable to use hACE2, we modeled the RBD domain of the sequences from Uganda (PDF-2370, PDF-2386) and Rwanda (PRD-0038). Unlike other non-ACE2 binders, homology modeling suggests that the RBDs of these viruses from Africa are structurally similar to SARS-CoV-1 and SARS-CoV-2 (Figure 4A). However, modeling the interaction with hACE2 reveals amino acid differences at key interfacial positions that can help explain the lack of interaction observed for the rVSV-Uganda and rVSV-Rwanda viruses (Figure 4B-C). There are four regions of the RBD that lie within 10Å of the interface with hACE2, one of which is the receptor binding ridge (SARS-CoV-1 residues 459-477) that is critical for hACE2 binding [32,40]. We have designated the remaining regions as regions 1 (residues 390-408), 2 (residues 426-443), and 3 (residues 478-491) (Figure 5).

**Figure 4.**
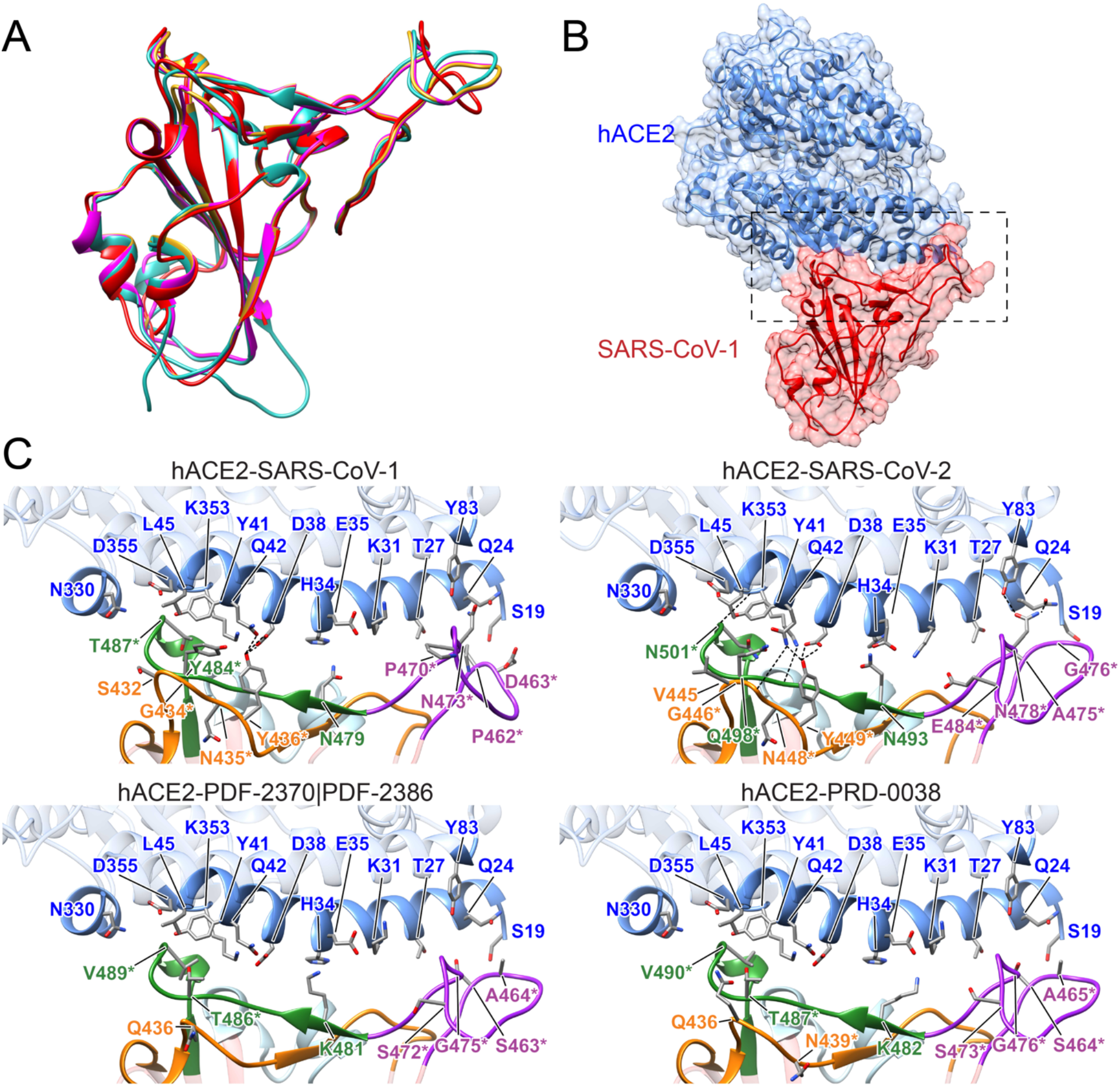
Structural modeling of sarbecovirus RBDs found in Uganda and Rwanda. (A) Structural superposition of the X-ray structures for the RBDs in SARS-CoV-1 (PDB 2ajf, red) [41] and SARS-CoV-2 (PDB 6m0j, cyan) [82] and homology models for SARS-CoV found in Uganda (PDF-2370 and PDF-2386, magenta) and Rwanda (PRD-0038, yellow). (B) Overview of the X-ray structure of SAR-CoV-1 RBD (red) bound to hACE2 (blue) (PDB 2ajf, red) [41]. (C) Close-up view of the interface between hACE2 (blue) and RBDs in SARS-CoV-1 (PDB 2ajf, top left) [41] and SARS-CoV-2 (PDB 6m0j, top right) [82] and homology models for viruses found in Uganda (PDF-2370 and PDF-2386, bottom, left) and Rwanda (PRD-0038, bottom, right). The color of the RBD loops corresponds to the colors of the labeled sequence regions in Figure 5: region 1 in cyan, region 2 in orange, the receptor binding ridge in purple, and region 3 in green. Labeled RBD residues correspond to interfacial residues whose identity differ in African sarbecoviruses and SARS-CoV-1 or SARS-CoV-2 (labels are included in all four panels to facilitate the identification of counterpart residues in each virus). Asterisks denote residues whose identity is not shared by any ACE-2 binding SARS-CoV as dictated by Figure 5. Labeled hACE2 residues correspond to residues within 5Å of RBD residues depicted.

**Figure 5:**
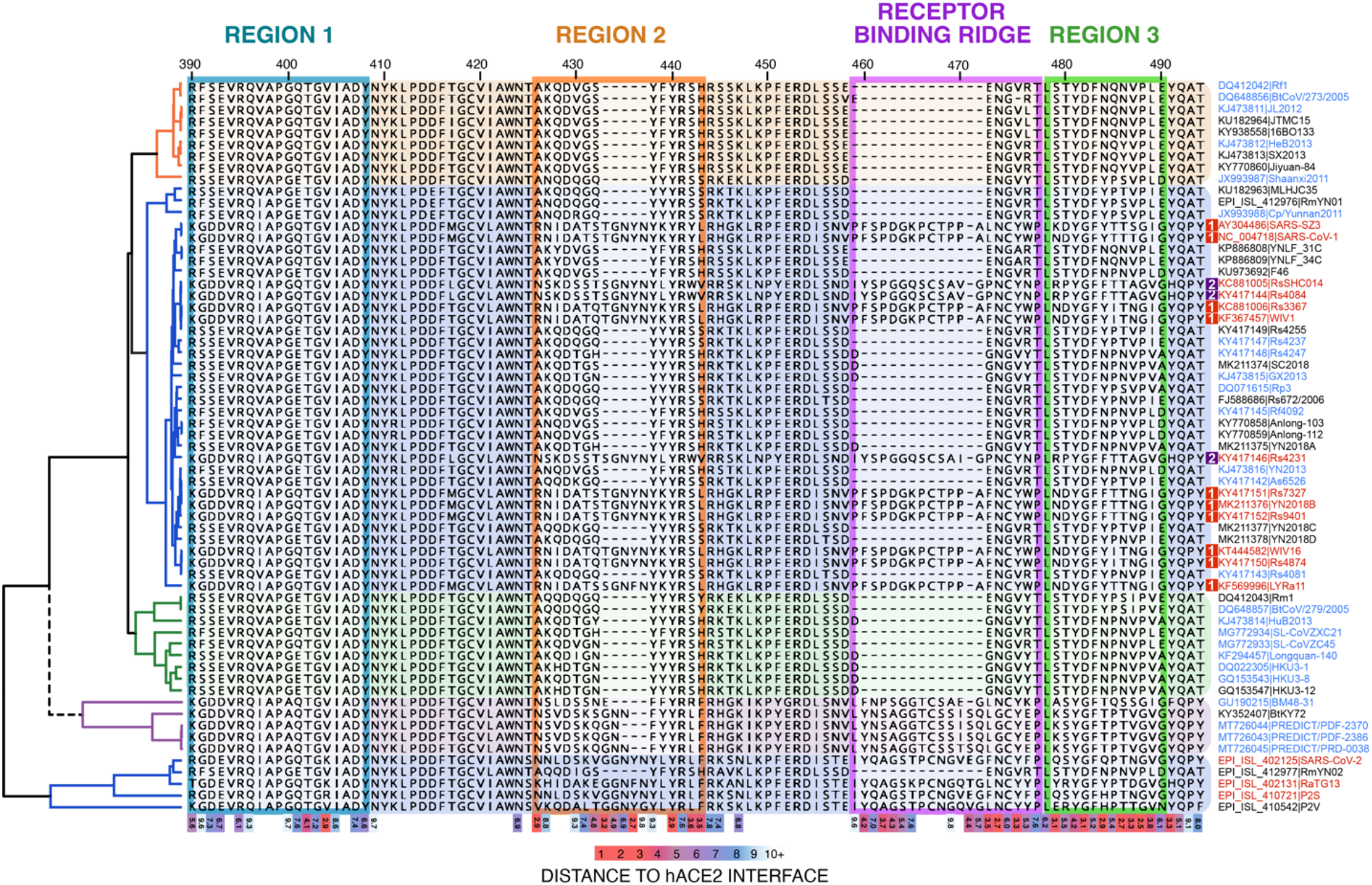
The phylogenetic backbone of the RdRp gene alongside the amino acid sequences of the RBM. Amino acid numbering is relative to SARS-CoV-1. Virus names in red font are known hACE2 users, those in blue are known non-users, and those in black have not been tested. Residues within 10Å of the interface with hACE2 are considered interfacial, and exact distances between each interfacial residue and the closest hACE2 residue (based on structural modeling of SARS-CoV-1 bound with hACE2) are shown along the bottom. Residues that are closer to the interface (3Å or less) and thus make strong interactions with hACE2 are shown in red, and as distance increases this color transitions to purple, blue, and finally to white. The receptor binding ridge sequences are highlighted in purple and the remaining interfacial segments have been numbered regions 1, 2, and 3 for clarity within the main text. The colors of these regions correspond with the colors in the structural models of Figure 4. The branch leading to Lineage 4 is dashed to demonstrate uncertainty in its positioning.

The sarbecoviruses from Africa evaluated here have a 2-3 amino acid deletion (SARS-CoV-1 residues 434-436) in region 2 (Figure 5). As many of the residues in this region make close contact with hACE2 (<5Å), it is possible that this contributes to the disruption of hACE2 binding. One of these residues, Y436, establishes hydrogen bonds with human ACE residues D38 and Q42 in both SARS-CoV-1 and SARS-CoV2 (Figure 4C). Notably, all other non-ACE2 binders also have deletions in residues 432-436. While this deletion is thought to interfere or reduce binding, restoring a similar deletion (SARS-CoV-1 residues 432-437) in the S protein of a European CoV (BM48-31) with the corresponding consensus segment obtained from Lineage 1 ACE2-binding viruses did not restore hACE2-mediated entry; only replacing the receptor-binding motif (RBM) increased hACE2-mediated entry [8].

Moreover, sarbecoviruses from Africa contain additional amino acid changes at the interface that can also contribute to hACE2 binding disruption (Figure 4C). hACE2 contains two hotspots (K31 and K353) that are crucial targets for binding by SARS-RBDs and amino acid variations in the RBD sequence enclosing these ACE2 hotspots have been shown to shape viral infectivity, pathogenesis, and determine the host range of SARS-CoV-1 [41–43]. All sarbecoviruses from Africa contain a Lys (K) at SARS-CoV-1 position 479 within region 3 (positions 481 and 482 for Uganda and Rwanda, respectively), which makes contact with these ACE2 hotspots (as compared to N479 or Q493 in SARS-CoV-1 and 2 respectively; Figure 4C). K479 decreases binding affinity by more than 20-fold in SARS-CoV-1 [44]. The negative contribution of K479 in region 3 is likely due to unfavorable electrostatic contributions with ACE2 hotspot K31 (Figure 4C) [42,45]. On the other hand, SARS-CoV-1 residue T487 (N501 in SARS-CoV-2) interacts with ACE2 hotspot K353 and has a Val (V) in the viruses from Africa (residues 489 and 490) (Figure 5). As with residue 479, the amino acid identity at position 487 contributes to the enhanced hACE2 binding observed in SARS-CoV-2 [42,43,45]. The presence of a hydrophobic residue at position 487, not previously observed in any ACE2 binding sarbecovirus, might lead to a local rearrangement at the K353 hotspot that hinders hACE2 binding. Indeed, most non-ACE2 binders have a Val (V) in SARS-CoV-1 position 487 (Figure 5).

Finally, the receptor binding ridge, which is conspicuously absent from all non-ACE2 binders, is present in the sarbecoviruses from Africa but has amino acid variations that differ significantly from both SARS-CoV-1 and SARS-CoV-2 (Figure 5). Changes in the structure of this ridge contribute to increased binding affinity of SARS-CoV-2, as a Pro-Pro-Ala (PPA) motif in SARS-CoV-1 (residues 469-471) replaced with Gly-Val-Glu-Gly (GVEG) in SARS-CoV-2 results in a more compact loop and better binding with hACE2 [32]. Changes within this ridge may be negatively contributing to hACE2 binding of viruses from Africa, which have Ser-Thr-Ser-Gln (STSQ) or Ser-Iso-Ser-Gln (SISQ) in this position (Figure 4C and 5).

While our studies suggest that these viruses from Africa do not utilize hACE2, it is not clear whether they are still ACE2-users but are adapted to divergent forms of bat ACE2 in their natural hosts. The specific bat host species for the Uganda and Rwanda viruses reported here could not be definitively identified in the field or in the lab, but are all genetically identical. They may represent a cryptic species, as the mitochondrial sequences are ~94% identical with *Rhinolophus ferrumequinum* in the cytochrome oxidase I gene (COI) and ~96% identical with *Rhinolophus clivosus* in the cytochrome b (cytb) gene, each of which have been deposited in GenBank (accessions MT738926-MT738928, MT732776). We were also able to extract ACE2 sequences from the deep sequencing reads of PDF-2370 (GenBank accession MW183243) to compare it to ACE2 sequences from species that are known to host ACE2 binders (human, civet, pangolin), non-ACE2 binders (*R. macrotis, pearsonii, pusillus, ferrumequinum*), and both (*R. sinicus*). Comparison of the ACE2 sequences shows that they are highly similar, with only a few amino acids that are changed in hosts of viruses that utilize ACE2 compared to the host of our African bat sample (Supplementary File 1). *R. sinicus* in particular is a known host of viruses that utilize ACE2 as well as viruses with the deletions that do not, suggesting that adaptation to divergent bat ACE2 is not a likely explanation for the deviation in sequence and structure of the RBD of viruses with deletions, including the novel sarbecoviruses from Uganda and Rwanda. These findings provide additional structural evidence that aids in distinguishing viruses which bind ACE2 from those that do not. They also demonstrate that ACE2 usage within sarbecoviruses is restricted to those viruses within the SARS-CoV-1 and SARS-CoV-2 clade in the RBD (Lineages 1 and 5, Figure 2).

The finding of discordant evolutionary trees for RdRp and the RBD in Figure 2 more strongly supports a recombination scenario; however, to consider an alternate scenario where ACE2 usage arose in Lineages 1 and 5 independently through convergent evolution, we compared the RdRp phylogeny with the amino acid sequences of the interfacial residues in the RBD (Figure 5). When mapped to the RdRp tree, the ‘extra’ RBD sequence present in the ACE2-using viruses is conspicuous within the Lineage 1 clade of otherwise non-ACE2-using viruses that have large deletions. We also note that there are two distinct groups of RBD sequences within ACE2-using Lineage 1 viruses: Type 1, containing SARS-CoV-1, SARS-SZ3 (civet), Rs3367, WIV1, Rs7327, YN2018B, Rs9401, WIV16, Rs4874, and LYRa11, and Type 2, containing Rs4231, Rs4084, and RsSHC014. Further, RmYN02 is within the Lineage 5 clade of ACE2-using viruses in RdRp but its RBD sequence contains both deletions (Figure 5). Without recombination, the viruses with deletions in region 2 and in the receptor binding ridge would have had to be gained and lost in precisely the same positions for ACE2-using Lineage 1 viruses and RmYN02, respectively, which is not a parsimonious explanation. The phylogeny and sequence in Figure 5 also illustrate that ACE2-usage appears to be an ancestral trait conserved in Lineage 5 [39] and a derived trait in each of the 13 Lineage 1 viruses with ACE2-using structure.

Finally, we further investigated support for the recombination scenario by examining the region of sequence between RdRp and the RBD for possible breakpoints. Only the 13 Lineage 1 viruses with ACE2-using structure were targets of this analysis as we were primarily interested in explaining the discordant phylogeny and variation in ACE2 usage (Figure 2), not in fully describing the recombination history of every sarbecovirus. Using 3SEQ, we show that all of the ACE2-using Lineage 1 sequences show extensive evidence of recombination within S1 and the RBD specifically (Table 2, Figure 6A). Further, the assignment of the parental sequence that donated the recombinant region (the minor parent) always resulted in the identification of one of the other recombinant sequences. This would not have been possible, as the recombinant region would have had to come from somewhere other than these 13 sequences, indicating that the true minor parent does not exist in our alignment. Using these breakpoints, we designated six subregions that were relatively free of recombination within these 13 sequences, mirroring the approach of Boni et al. 2020 [39], and built phylogenetic trees for each region. We show that in orf1ab (region A) and S2 (region F) these 13 sequences fall within Lineage 1, but within S1 and particularly the RBD (B through E) they switch phylogenetic positions and cluster with Lineage 5 (Figure 6B), supporting the recombination scenario.

**Figure 6.**
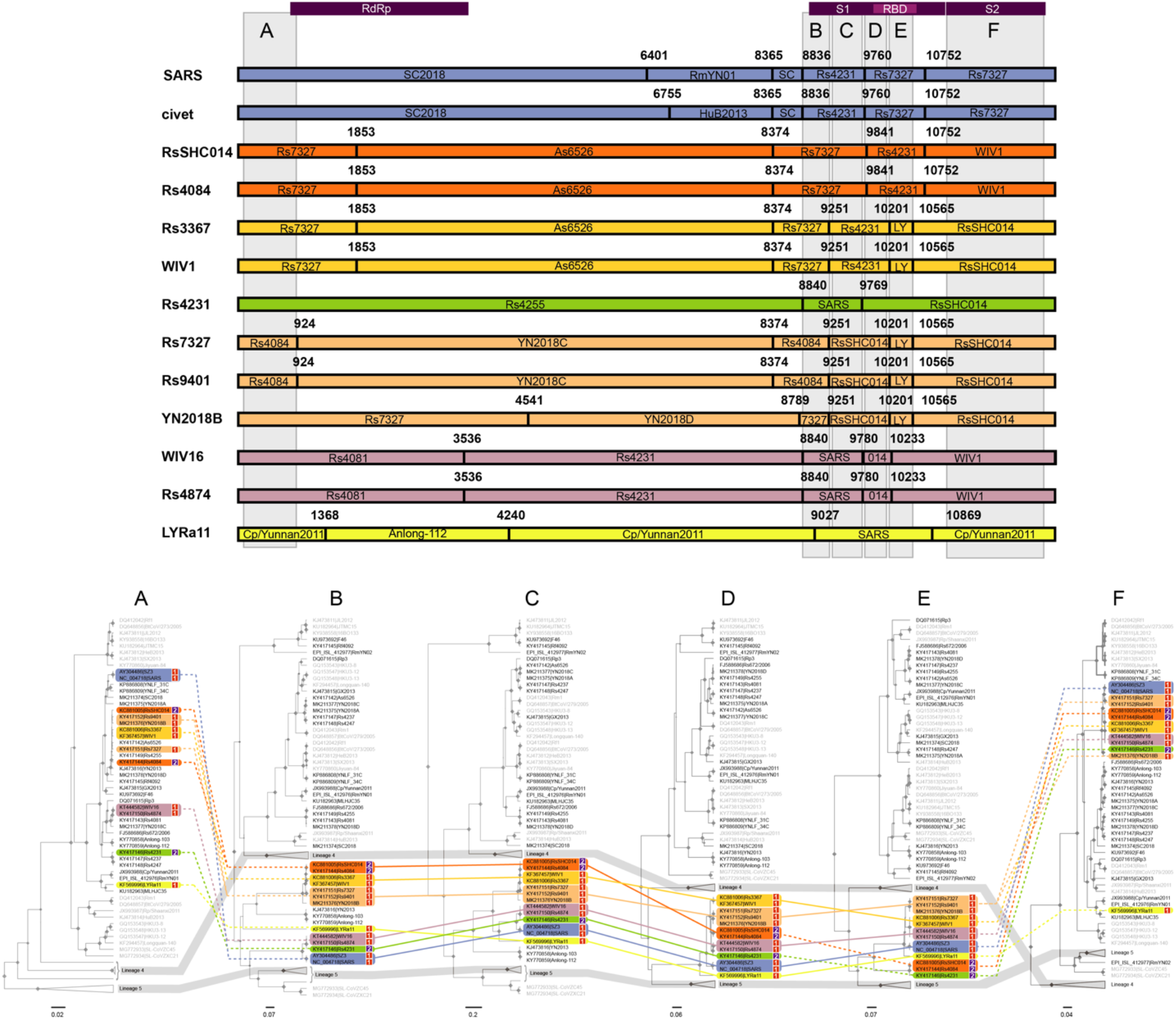
Recombination breakpoints detected in Lineage 1 ACE2-using sequences. The top of this figure illustrates that the recombination suggested by the change in topology in Figure 2 for 13 Lineage 1 viruses is supported by formal breakpoint analysis. The breakpoints detected for each of the 13 recombinant Lineage 1 sequences with ACE2-using structure (no deletions) are shown. Sequences that are nearly identical are colored the same for simplicity. The bars represent the sequence of genome beginning 750 bp before RdRp spanning through the end of S2 (SARS-CoV-2 nucleotides 12,681 through 25,176) and each box within represents a recombinant section within the sequence. The breakpoints correspond to those identified in Table 2. Numbering is relative to the alignment. The parental sequence is shown within each box. Sequences identified as the minor parent by 3SEQ were labeled within the breakpoint margins and the major parent outside. Six regions where these sequences appear to be free of recombination are labeled A-F and a corresponding phylogeny for each region is shown below. Regions A and E were further tested for recombination breakpoints in all sequences, not just the 13 Lineage 1 viruses, and were found to be breakpoint-free. The topology of regions A and E is not different enough from Figure 2 to suggest that recombination within RdRp or RBD significantly changed the interpretation of our results. For each region, sequences were tracked with connecting lines of corresponding color to identify where recombination may have occurred between Lineage 1 and Lineage 5 and hypothesized events are specifically marked with dotted lines. This highlights the secondary recombination of Rs4084 and RsSHC014 in region E on top of the primary recombination in regions B through E. Sequence names of Lineage 2 and 3 viruses are greyed out and Lineages 4 and 5 are collapsed and highlighted in darker grey to make the changes in topology between the trees more visible.

Despite only investigating the Lineage 1 recombinants for the locations of sequence breakpoints, the phylogenetic trees provide evidence that recombination has occurred frequently in other sarbecoviruses in this genomic region as well (Figure 6B). Of note, Rs4084 and RsSHC014 cluster with Type 1 RBDs in regions B, C, and D, but with swap to cluster with Rs4231 (Type 2) in Region E, even though Rs4084, RsSHC014, WIV1, and Rs3367 are all nearly identical in every other region. This suggests that a WIV1/Rs3367-like Type 1 virus which had already undergone recombination in regions B through E underwent a second recombination event with a Type 2 virus on top of the first in region E. A number of other viruses also appear to have recombinant history in regions B, C, and D (SL-CoVZC45 and SL-CoVZXC21, YN2013, Anlong-103, and Anlong 112), but these viruses do not show evidence of recombination that spans the RBD in region E, which contains the amino acid deletions in region 2 and the receptor binding ridge and appears to primarily determine ACE2-using potential. The frequency of recombination in this region among Lineage 1 viruses strongly supports the hypothesis that after ACE2-usage was acquired in Lineage 1, it subsequently spread throughout the clade via additional recombination events with other Lineage 1 viruses.

As all of our evidence supports a recombination scenario over convergent evolution, we sought to construct a possible timeline of events that could explain our observations. Using tip dating in BEAST2, we constructed a time-calibrated phylogeny for RdRp using a substitution rate prior inferred from Boni et al. 2020 [39]. Using the RdRp tree as an evolutionary backbone, the deletions in region 2 and the receptor binding ridge of the RBD appear to have been lost in a stepwise fashion (Figure 5). The small deletion in region 2 likely arose first, before the diversification of Lineage 4 in Africa and Europe (Figure 5) and was dated using the MRCA of Lineages 1, 2, 3 and 4 (Figure 8). Alternatively, as the boundaries of the deletion in region 2 in Lineage 4 and Lineages 1, 2, and 3 do not align perfectly and there is uncertainty in the position of this branch in the phylogeny, it is equally possible that this deletion was lost independently in Lineage 4. The larger deletion in the receptor binding ridge, not present in known sequences from Lineage 4, likely arose second, but before the diversification of Lineages 1, 2, and 3 (Figure 5) and was dated with the MRCA of these three lineages (Figure 8). Because no ACE2-using viruses have been discovered in Lineage 2 or 3 to date, we propose that the re-appearance of this trait arose after the MRCA of Lineage 1 on the tree (Figure 8). As SARS-CoV-1 was the earliest Lineage 1 virus sequenced with ACE2-using structure, the emergence of ACE2 usage in Lineage 1 must have occurred in the time between the MRCA of Lineage 1 (1852, 95% HPD 1804-1901) and the emergence of SARS-CoV-1 in 2003.

Next, we constructed a time-calibrated phylogeny for RBD with a strict MRCA age prior informed by the estimation of the tree height in RdRp (see *Methods*), such that the timescale would be comparable even though the evolutionary rates between these two regions likely are not the same (Figure 7). To account for variability in lineage-specific substitution rates, we also generated a time-calibrated model using a relaxed lognormal clock (Figure 7). Comparing the time-calibrated RBD tree to the time-calibrated RdRp tree, the divergence dates for the two types of RBD sequence observed in the recombinant Lineage 1 sequences are incompatible, suggesting that more than one recombination event donating ACE2 usage from Lineage 5 to Lineage 1 must have occurred. The 13 Lineage 1 recombinants (both Type 1 and Type 2) coalesce between 119-216 years ago in RdRp and between 259-490 years ago in the RBD (Figure 7). If these time estimates reflect true rates of diversification, a single introduction of the ACE2-using phenotype via recombination would not allow enough time for the sequence divergence between Type 1 and Type 2 RBDs to accumulate, even when accounting for the substitution rate in RBD being estimated as an order of magnitude higher than that of RdRp (5.248e-4 in RdRp, 2.181e-3 in RBD). Further, the substitution rate that would be needed for the observed sequence divergence in the RBD of the 13 recombinants to have accumulated since their MRCA in RdRp (1852) is more than double the estimated rate of our time-calibrated tree (5.899e-3). Even with a relaxed clock assumption, the maximum value of the posterior distribution of the mean rate is only 4.733e-3. From this, we conclude that two independent recombination events occurred between Lineage 5 and Lineage 1 resulting in two distinct RBD types.

**Figure 7.**
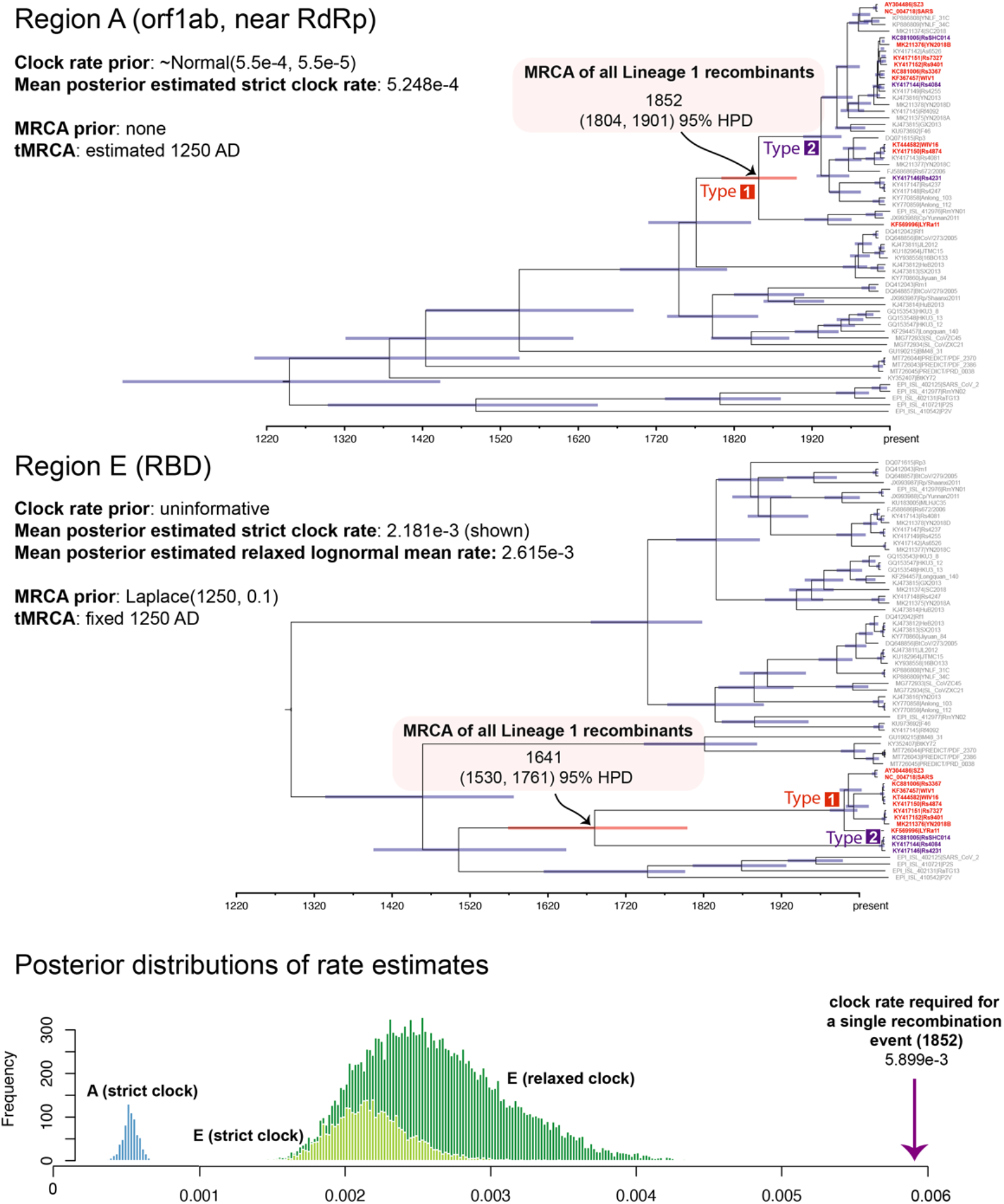
Time-calibrated phylogenies for recombination-free regions of the genome. Breakpoint-free regions A and E from Figure 6 were chosen for time calibration since evidence of recombination was found in both RdRp and RBD. Both regions A and E were free of recombination for all sequences included in the tree, ensuring the best possible dating estimates. The MRCA of all Lineage 1 recombinants and its corresponding divergence date are labeled on each tree, demonstrating that the MRCA in region E (within the RBD) is much older than the MRCA in region A (proxy for RdRp, see Figure 6). This suggests that there would not have been enough time for the RBDs of the recombinants to diversify to the extent shown here if only a single recombination event occurred between Lineage 5 and Lineage 1. The MRCAs of each type are labeled in red (Type 1) and purple (Type 2). Posterior distributions of rate estimates are also shown for each model as well as for a relaxed clock model of region E. For the observed sequence divergence in region E to have accumulated since the MRCA of the 13 recombinants in region A (1852), a clock rate of 5.899e-3 would be required, which is well outside the posterior distributions estimated by both our strict and relaxed clock models.

We propose two main hypotheses for the acquisition and spread of the two distinct RBD types donating ACE2 usage from Lineage 5 to Lineage 1. The recombination hypothesis posits that two recombination events donated Type 1 and Type 2 RBD sequence from Lineage 5 to Lineage 1; however, these two events are insufficient to explain the non-monophyletic pattern of ACE2 usage in Lineage 1. We further hypothesize that whichever Lineage 1 virus first gained Type 1 and Type 2 ACE2 usage in each group then donated the trait to other Lineage 1 viruses through subsequent recombination events (Figure 8). It is difficult to approximate a date for such an event, but the MRCA of the Type 1 recombinants in the RBD may be a close estimation (between 42 and 77 years ago) (Figure 7). The events must have been recent enough that the observed diversity of Type 2 RBD sequences is quite low, yet not so recent such that there would not have been time for recombination to have occurred twice in region E for sequences Rs4084 and RsSHC014 (Figure 6B).

**Figure 8.**
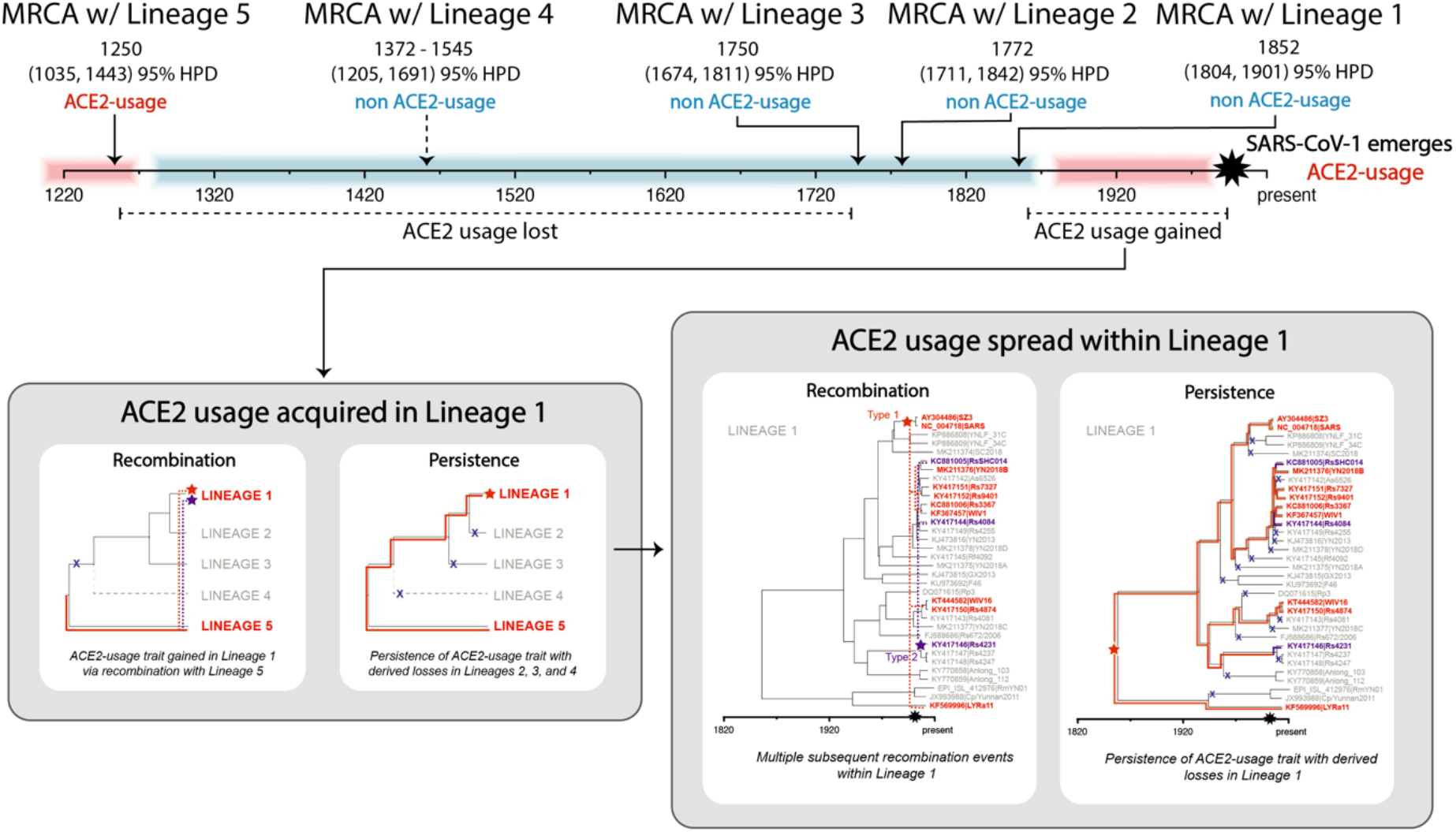
Proposed timeline of deletion and recombination events. The timeline demonstrates the sequence of events that led to loss of ACE2 usage in Lineages 2, 3, and 4 and gain of ACE2 usage within Lineage 1, leading to the emergence of SARS-CoV-1. Events are dated with MRCA age estimates; however, the exact intention is less to provide exact dates and more to suggest a particular order of events, which is strongly supported by the posterior probabilities of the time-calibrated phylogenies. The arrow for the Lineage 4 event is again dashed to demonstrate uncertainty in its positioning. We illustrate two hypotheses for the acquisition and subsequent spread of ACE2 usage in Lineage 1: recombination and persistence. The recombination hypothesis is much more parsimonious, as persistence would require multiple independent deletion events to generate the observed pattern of ACE2 usage.

The second hypothesis and only remaining possibility for ACE2 usage in Lineage 1 (besides convergence) is that perhaps the trait persisted in this Lineage from the ancestral state (Figure 8). Because no viruses demonstrating ACE2 usage have been discovered in Lineages 2, 3, and 4, this would mean that the ACE2 usage trait would have been lost via deletion in these lineages. Further, because of the non-monophyletic branching order of these lineages, this would require multiple independent and identical losses of the region 2 and receptor binding ridge deletions in all three of these lineages. If this did indeed occur, in order to then observe the pattern of ACE2 usage in Lineage 1 where some viruses, but not all, have the ACE2 usage trait, further independent losses would be required in individual viruses. In much the same manner as convergence would require multiple independent and identical events, persistence of ACE2 usage with multiple independent deletions for the entire clades of Lineages 2, 3, and 4 and only some of the viruses in Lineage 1 is also highly non-parsimonious. Persistence is also a poor explanation for the pattern of the two RBD types observed, particularly for Type 2, where the RBD sequences are highly similar but the RdRp sequences are quite divergent. If both genes were vertically inherited via persistence, we would expect these genes to have approximately equal MRCA ages. Instead, we observe that the MRCA age for Type 2 RBDs in region E are much younger than for RdRp.

## Discussion

### ACE2 usage in Lineage 1 viruses was acquired via recombination

At first glance, ACE2 usage does not appear to be phylogenetically conserved among sarbecoviruses, especially since many phylogenies are built using RdRp. This naturally leads to the hypothesis that ACE2 usage arose independently in SARS-CoV-1 and SARS-CoV-2 via convergent evolution. This has been suggested previously for another ACE2-using human coronavirus, NL63 [46]. However, a phylogeny constructed using the RBD perfectly separates viruses that have been shown to utilize ACE2 from those that do not (Figure 2). Viruses that cannot utilize ACE2 have significant differences in their RBDs, including large deletions in critical interfacial residues and low amino acid identity with viruses that do use ACE2 (Figure 5). Notably, in addition to the large deletions, viruses that cannot use ACE2 deviate considerably at the interacting surface, including positions that play fundamental roles dictating binding and cross-species transmission [32,41,44,47]. It is unknown whether viruses that cannot use hACE2 are utilizing bat ACE2 or an entirely different receptor altogether, but since mammalian ACE2 is so conserved [48,49] and ACE2-using viruses demonstrate broad host tropism [42,50–52], we hypothesize that there is likely a different receptor involved for the non-ACE2 users (see Supplementary File 1).

The difference in topology, specifically in the positioning of ACE2-using Lineage 1 viruses, between RdRp and RBD trees suggests that the ability to use ACE2 was introduced into Lineage 1 by recombination between a recent ancestor of the ACE2-using Lineage 1 viruses (including SARS-CoV-1) and an undiscovered Lineage 5 virus in the RBD. As there are two types of closely related RBD sequences in the recombinant Lineage 1 viruses (Figure 2) with incompatible divergence dates (Figure 7), we suggest that two such recombination events occurred between Lineage 1 and Lineage 5 (Figure 8) independently introducing ACE2-usage into Lineage 1. The non-monophyletic nature of ACE2 usage within Lineage 1 can then be most parsimoniously explained by secondary intra-lineage recombination events (Figure 8). It is possible that both hypotheses are partially true and that both intra-lineage recombination as well as the persistence of this trait alongside sister Lineage 1 viruses without the trait gave rise to the observed patterns of Type 1 and Type 2 ACE2 usage within Lineage 1. It is also very possible that further sampling may illuminate that some of the events proposed here have been distorted by sampling bias. We have estimated that these events may have occurred roughly within the last two centuries, though this estimate will likely change with further sampling as well. Our intention is not necessarily to date these events exactly, but rather to infer their order relative to each other and to make hypotheses based on this order of events. Confidence intervals for many node dates overlap, but high posterior probabilities on internal nodes indicate that events most likely occurred in a certain order.

Our conclusion that ACE2 usage originated in Lineage 5 and was introduced into Lineage 1 by recombination is based on phylogenetics; however, studies of recombination using phylogenetics are often limited in their ability to definitively determine the direction of recombination. Nonetheless, there are several lines of evidence that support the direction having occurred from Lineage 5 to Lineage 1. First, recombination is notoriously more frequent in spike compared to orf1ab [39,53,54]. Second, Lineage 5 constitutes the base of the tree and has the oldest MRCA, meaning it likely shares more ancestral traits with the MRCA of all sarbecoviruses. Third, phylogenetic topology in orf1ab before the recombinant region of the genome mirrors that of S2 after the recombinant region (Figure 6A), orienting orf1ab/S2 as sequence from the major parent of the recombination event. And finally, that spike is the recombinant region as opposed to RdRp is also supported by numerous studies that have provided evidence that SARS-CoV-1 is recombinant and SARS-CoV-2 is not [3,13,15,55].

In order for recombination to have occurred between Lineage 1 and Lineage 5, these viruses must have had the opportunity to coinfect the same host cell. We demonstrate that recombination is possible given that viruses related to SARS-CoV-1 and −2 appear to share both geographic and host space in southwestern China and in *R. sinicus* and *R. affinis* bats. Highlighting that this previously known recombination event (i.e. SARS-CoV-1) occurred with a previously unknown group of viruses that are related to SARS-CoV-2 is an important finding of this study and demonstrates that recombination is an important driver of spillover for sarbecoviruses.

### A series of deletion events most likely resulted in the ancestral loss of ACE2 usage in Lineages 1-4

Using the RdRp tree as the evolutionary history to which to compare because of its stability and relative lack of recombination, sequences without the deletions in the RBD most likely represent the ancestral state, as the SARS-CoV-2 Lineage 5 viruses at the base of the tree do not show this trait (Figure 2). This is in accordance with the findings of Boni et al. [39]. Alternatively, it is possible that the deletion state is the ancestral state, and that this ancestral deletion state was conserved in Lineages 1, 2, and 3; however, insertions acquired during the evolution of Lineages 4 and 5 would have had to have occurred independently, which is less parsimonious. Persistence of the ACE2 usage trait from the MRCA of Lineage 5 all the way to Lineage 1 is also not parsimonious, as the RBD deletions would have had to have been lost many times independently (Figure 8).

Further, the viruses from bats in Africa and Europe have one of the two deletions, which may indicate that these are descendant from an evolutionary intermediate and support a stepwise deletion hypothesis; however, this hypothesis hinges completely on the uncertain positioning of Lineage 4 on the phylogeny, which may support independent deletion within region 2 in Lineage 4 instead. Since ACE2-using Lineage 1 viruses including SARS-CoV-1 are nested within a clade of viruses that all have both deletions, this implies that both deletions arose before the diversification of Lineages 1, 2, and 3 viruses (Figures 5 and 8). According to the branching order shown here, the smaller deletion in region 2 was likely acquired earliest, before the diversification of the clades into Africa and Europe, since it is shared by all clades with the exception of SARS-CoV-2 Lineage 5 at the base of the tree (Figure 5). These large deletions in the RBD-ACE2 interface and the similarity of Rhinolophid and hACE2 also suggest that non-ACE2-using viruses, including Lineages 1, 2, 3, and 4, are using at least one receptor other than ACE2 [8,36].

### ACE2 usage is not well explained by convergent evolution

Under a hypothetical convergent evolution scenario, large insertions would have had to be reacquired in precisely the same regions from which they were lost within the RBD independently in ACE2-using Lineage 1 viruses. The most parsimonious argument is that ACE2-using Lineage 1 viruses are descendent from at least two recombinant viruses (containing Types 1 and 2 RBDs) and that recombination best explains the non-monophyletic pattern of ACE2 usage within the *Sarbecovirus* subgenus. In contrast, human coronavirus NL63 is an alphacoronavirus that is also a hACE2 user but most likely represents a true case of convergent evolution. The RBD of SARS-CoV-1 and SARS-CoV-2 are structurally identical, while NL63 has a different structural fold, suggesting that they are not evolutionarily homologous [46]. Nonetheless, NL63 also binds to hACE2 in the same region – suggesting all of the ACE2-using viruses have converged towards this interaction mode [46].

Additional evidence supports a recombination scenario over convergent evolution, including (i) the detection of statistically supported recombination breakpoints in all ACE2-using Lineage 1 viruses between RdRp and the RBD, and (ii) a growing number of reports identifying recombination in the spike gene of other CoVs [22,56–59]. We also highlight an additional unreported recombination event between Lineage 5 and Lineage 1 giving rise to RmYN02 that further demonstrates the importance of this evolutionary mechanism. We observed that the Lineage 5 bat virus RmYN02, which is highly similar to SARS-CoV-2 within the RdRp, actually has a RBD with the Lineage 1 deletion trait associated with the inability to use ACE2. This indicates a recombination in the opposite direction, From Lineage 1 to Lineage 5, and is again consistent with their overlapping host and geographic ranges. The RmYN02 virus was sequenced from a pooled sample that also contained a second strain, RmYN01, so the possibility that the assembled RmYN02 sequence is chimeric cannot be ruled out. However, both RmYN01 and RmYN02 have deletions in the RBD, so whether or not the sequence is chimeric, it is most likely still recombinant. Again, recombination is a much more parsimonious explanation for the loss of ACE2 usage in RmYN02 rather than convergence, which would require independent and identical deletions in the interfacial residues of the RBD.

### Differences in receptor usage within sarbecoviruses would explain observed phylogeographic patterns

Lineage 1 and Lineage 5 viruses appear to occupy the same geographic space, which is necessary for the opportunity to recombine to exist. However, the co-circulation of these distantly phylogenetically related viruses is a notable deviation from previous observations that show sarbecovirus phylogeny mirrors geography. It is unknown why Lineages 1-4 show strong phylogeographic clustering. Isolation by distance (IBD) is one ecological mechanism that could explain concordance between phylogeny and geography; however, this would not explain why Lineage 5 deviates from this pattern and overlaps geographically with Lineage 1. Instead, we hypothesize that immune cross-reactivity between closely related viruses within hosts results in indirect competitive exclusion and priority effects, and that this explains the phylogeographic signal of Lineages 1-3. Antibodies against the spike protein are critical components of the immune response against CoVs [60–62]. Hosts that have been infected by one sarbecovirus may be immunologically resistant to infection from a related sarbecovirus, leading to geographic exclusion of closely related strains and a pattern of evolution that is concordant with geography despite the fact that species and individuals are not strictly confined (Figure 1). It is unlikely that this pattern is caused by differing competencies amongst *Rhinolophus* bats, as host-switching of these viruses appears to be common. The co-circulation of Lineage 5 viruses (including SARS-CoV-2 and related viruses) in the same species and the same geographic location as Lineage 1 viruses may suggest a release in the competitive interactions maintaining geographic specificity. This would preclude recognition by cross-reactive antibodies, such as those produced against the spike protein, and may be evolutionarily advantageous for the recombinant virus. Furthermore, if these two groups of viruses utilize different receptors, antibodies against one would be ineffective at excluding the other, potentially allowing both viral groups to infect the same hosts. If competitive release has indeed occurred among these viruses, it is likely that the SARS-CoV-2 clade is potentially much more diverse and geographically widespread than currently understood.

### Implications for future research

Here, we highlight the critical need for further surveillance specifically in southwestern China and surrounding regions in southeast Asia given that all ACE2-using bat viruses discovered to date were isolated from bats in Yunnan Province. If this holds true, it would support the hypothesis that SARS-CoV-2 originated in Yunnan or the surrounding regions of southwest China before the initial epidemic then amplified in Wuhan. Southeast Asia and parts of Europe and Africa have been previously identified as hotspots for sarbecoviruses [63], but increased surveillance will help characterize the true range of ACE2-using sarbecoviruses in particular. The receptors for viruses from northern China and other regions such as Europe and Africa remain unknown, and may not pose a threat to human health if they cannot utilize hACE2, though their potential to acquire hACE2-usage by recombination should be considered along with the potential for their existing spike proteins to use other human receptors for cell entry. It is unclear whether the lack of hACE2 binding for sarbecoviruses from Uganda and Rwanda is due to the small deletion in region 2 or to the numerous amino acid changes in other interfacial residues. It is possible that sarbecoviruses in Africa with different residues in these interfacial regions could potentially still use hACE2. It is also unknown whether the sarbecoviruses from Africa in particular use a different receptor altogether, or whether sarbecoviruses with the potential to utilize hACE2 without the region 2 deletion have also diversified into Africa or Europe. If competitive release between groups of viruses utilizing different receptors has indeed occurred, further surveillance is needed to determine the true extent of Lineage 5 viruses. In addition, experimental evidence to support or refute a competitive release hypothesis should be prioritized.

This study highlights that hACE2 usage is unpredictable using phylogenetic proximity to SARS-CoV-1 or SARS-CoV-2 in the RdRp gene. This is due to vastly different evolutionary histories in different parts of the viral genome due to recombination. Phylogenetic relatedness in the RdRp gene is not an appropriate proxy for pandemic potential among CoVs (the ‘nearest neighbor’ hypothesis). By extension, the consensus PCR assays most commonly used for surveillance and discovery, which mostly generate a small fragment of sequence from within this gene [64–66], are insufficient to predict hACE2 usage. Using phylogenetic distance in RdRp as a quantitative metric to predict the potential for emergence is tempting because of the large amount of data available, but this approach is unlikely to capture the biological underpinnings of emergence potential compared to more robust data sources such as full viral genome sequences. The current collection of full-length sarbecovirus genomes is heavily weighted toward China and *Rhinolophus* hosts, despite evidence of sarbecoviruses prevalent outside of China (such as in Africa) and in other mammalian hosts (such as pangolins). Further, investigations into determinants of pathogenicity and transmission for CoVs and the genomic signatures of such features will be an important step towards the prediction of viruses with spillover potential, and distinguishing those with pandemic potential.

Finally, these findings reiterate the importance of recombination as a driver of spillover and emergence, particularly in the spike gene. If SARS-CoV-1 gained the ability to use hACE2 through recombination, other non-ACE2-using viruses could become human health threats through recombination as well. We know that recombination occurs much more frequently than just this single event with SARS-CoV-1, as the RdRp phylogeny does not mirror host phylogeny and the RBD tree has significantly different topology across all geographic lineages. In addition, the bat virus RmYN02 appears to be recombinant in the opposite direction (Lineage 5 backbone with Lineage 1 RBD) [36], again supporting the hypothesis that recombination occurs between these lineages. Our analyses support two hypotheses: first, that sarbecoviruses frequently undergo recombination in this region of the genome, resulting in this pattern, and second, that sarbecoviruses are commonly shared amongst multiple host species, resulting in a lack of concordance with host species phylogeny and a reasonable opportunity for coinfection and recombination. Bats within the family *Rhinolophidae* have also repeatedly shown evidence of introgression between species [67–72], supporting the hypothesis that many species in this family have close contact with one another which may facilitate viral host switching. Given that we have shown that ACE2-using viruses are co-occurring with a large diversity of non-ACE2-using viruses in Yunnan Province and in a similar host landscape, recombination poses a significant threat to the emergence of novel sarbecoviruses [7].

With recombination constituting such an important variable in the emergence of novel CoVs, understanding the genetic and ecological determinants of this process is a critical avenue for future research. Here we have shown not only that recombination was involved in the emergence of SARS-CoV-1, but also demonstrated how knowledge of the evolutionary history of these viruses can be used to infer the potential for other viruses to spillover and emerge. Understanding this evolutionary process is highly dependent on factors influencing viral co-occurrence and recombination, such as the geographic range of these viruses and their bat hosts, competitive interactions with co-circulating viruses within the same hosts, and the range of host species these viruses are able to infect. Our understanding depends on the data we have available - the importance of generating more data for such investigations cannot be understated. Investing effort now into further sequencing these viruses and describing the mechanisms that underpin their circulation and capacity for spillover will have important payoffs for predicting and preventing sarbecovirus pandemics in the future.

## Methods

### Consensus PCR and sequencing of sarbecoviruses from Africa

Oral swabs, rectal swabs, whole blood, and urine samples collected from bats sampled and released in Uganda and Rwanda were assayed for CoVs using consensus PCR as previously described [22]. All sampling was conducted under UC Davis IACUC Protocol No. 16048. Bands of the expected size were purified and confirmed positive by Sanger sequencing and the PCR fragments were deposited to GenBank (accessions MT738926-MT738928, MT732776). Samples were subsequently deep sequenced using the Illumina HiSeq platform and reads were bioinformatically de novo assembled using MEGAHIT v1.2.8 [73] after quality control steps and subtraction of host reads using Bowtie2 v2.3.5. Contigs were aligned to a reference sequence and any overlaps or gaps were confirmed with iterative local alignment using Bowtie2. The full genome sequences are deposited in GenBank. Cytochrome b, cytochrome oxidase I. and ACE22 host sequences were also extracted bioinformatically where possible by mapping reads to *Rhinolophus ferrumequinum* reference genes using Bowtie2 and deposited in GenBank.

### Phylogenetic reconstruction

All publicly available full genome sarbecovirus sequences were collected from GenBank and SARS-CoV-2, pangolin virus genomes, RaTG13, and RmYN01/RmYN02 were downloaded from GISAID (Table 1). All relevant metadata (geographic origin, host species, date of collection) was retrieved from GenBank or the corresponding publications. The RdRp gene (nucleotides 13,431 to 16,222 based on SARS-CoV-2 sequence EPI_ISL_402125 from GISAID) and RBD region (nucleotides 22,506 to 23,174 based on the same SARS-CoV-2 reference genome) were extracted and aligned using Muscle v10.2.6. We chose RdRp as a backbone to which to compare because of the strong evolutionary constraints imposed by its fundamental biological role in viral replication [53]. Indeed, the RdRp is generally considered to be a primary genetic trait in viral taxonomy [1,38] and most viruses exhibit strong purifying selection in this gene [74]. Further, the orf1ab region of coronaviruses (which contains the RdRp) also tends to be more recombination-free as compared to the recombination-frequent latter half of the genome [39,54]. Since many of our conclusions are based around phylogenetic topology, we confirmed the robustness of the topology of our nucleotide trees by also building identical trees with alignments of other relatively stable genes in orf1ab frequently used for taxonomic classification [38] (Supplementary Figure S1). Phylogenetic reconstruction was performed using BEAST v2.6.3 [75] with partitioned codon positions, a GTR+Γ substitution model for each of the three codon positions, a constant size coalescent process prior, and a strict molecular clock model. Log files were examined using Tracer v1.7.1 to confirm that the model converged and that the effective sample size (ESS) for each parameter was at least 100. Chains were run until these convergence criteria were met (~2-10 million samples) and multiple chains were run independently to ensure convergence to the same estimates. Use of Beagle 2.1.2 was chosen to increase computational speed.

**Table 1.**
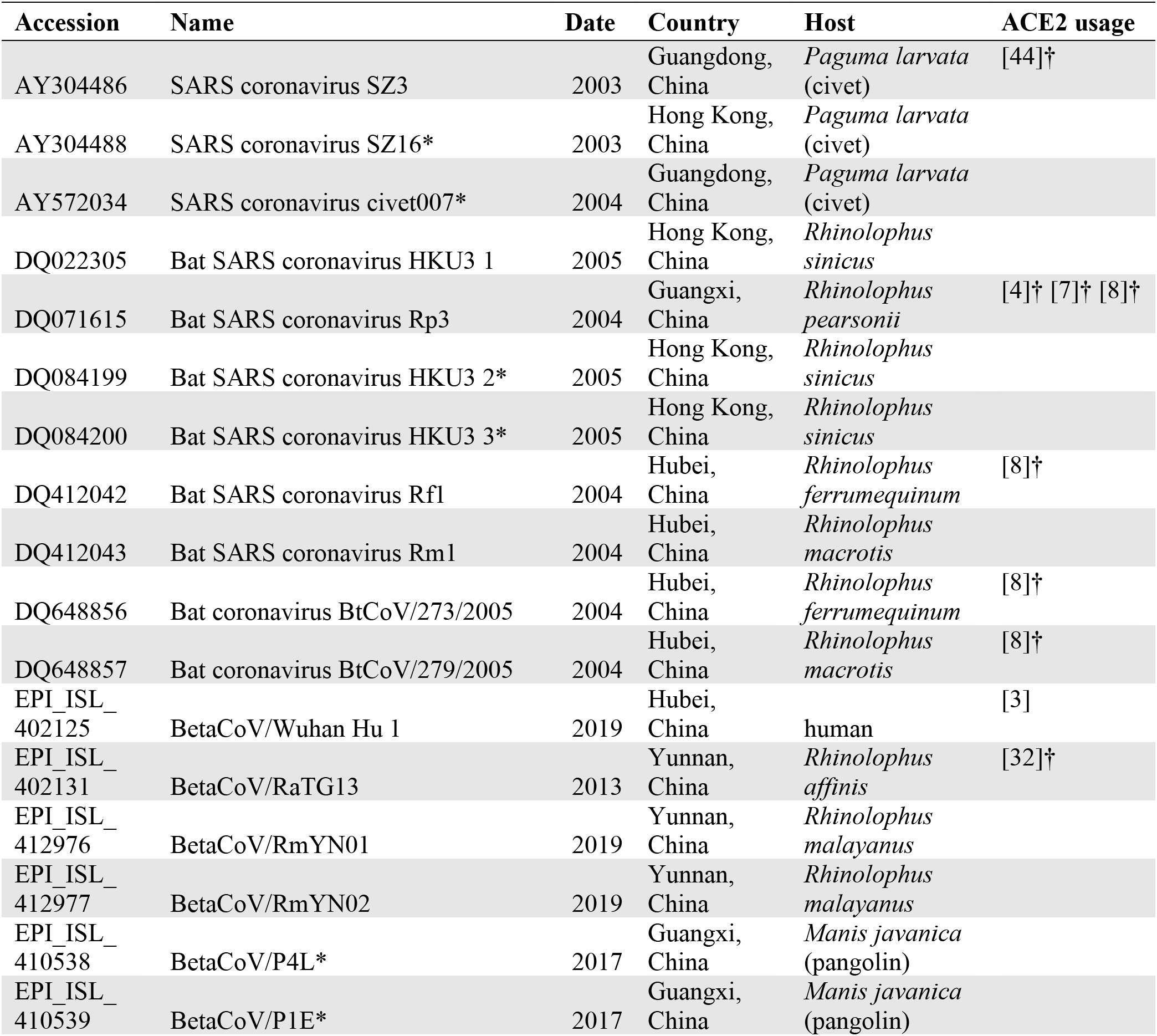
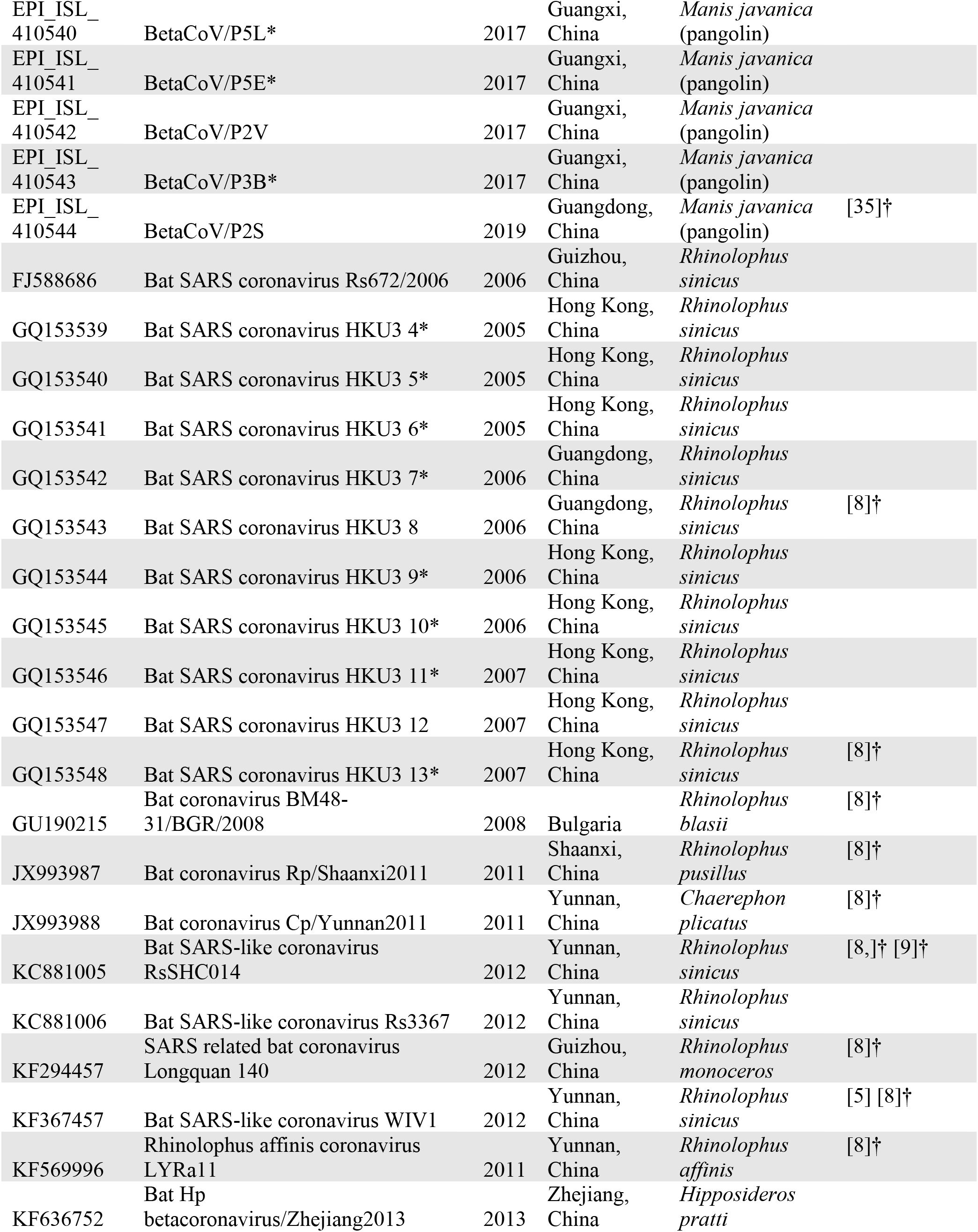
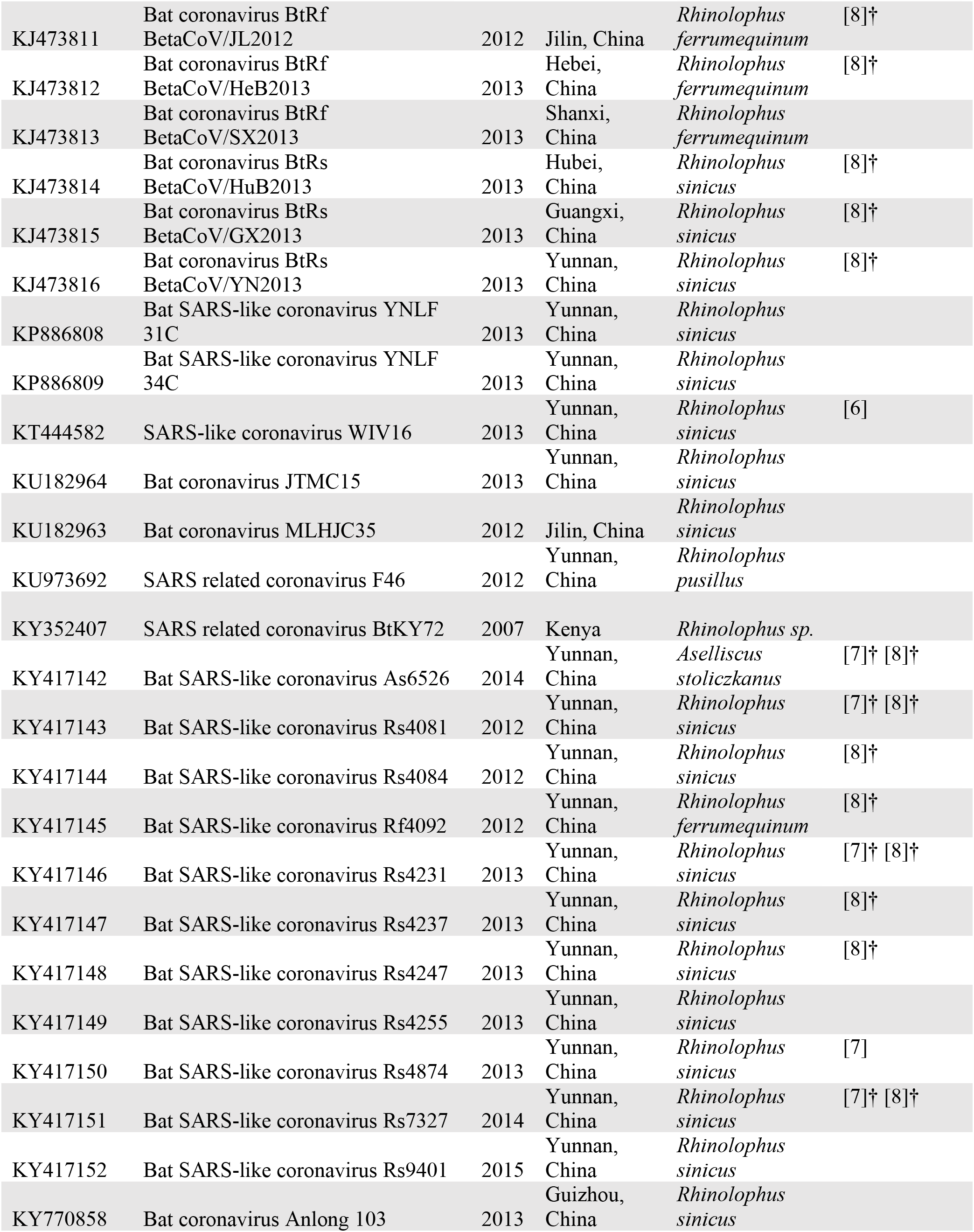
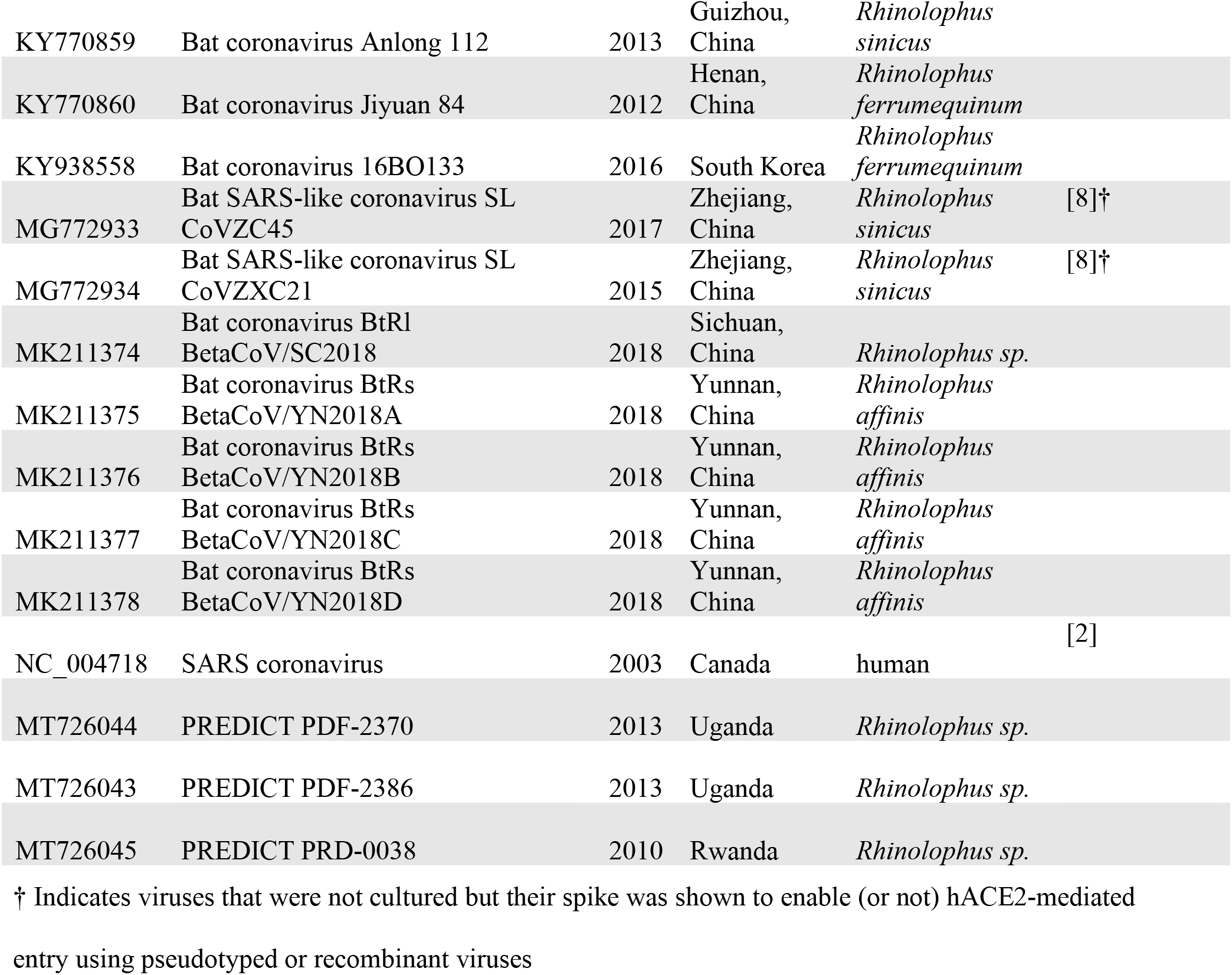
Full list of sequences and accession numbers used in this study. All accession numbers are from GenBank with the exception of those beginning with EPI_ISL, which are from GISAID. Metadata includes sequencing year, geographic origin, and host species. Sequence names marked with an asterisk (*) indicate those that were not included in the final phylogenetic reconstruction due to high genetic identity with another sequence in the alignment. Citations used to determine hACE2 binding capability are also included.

**Table 2.**
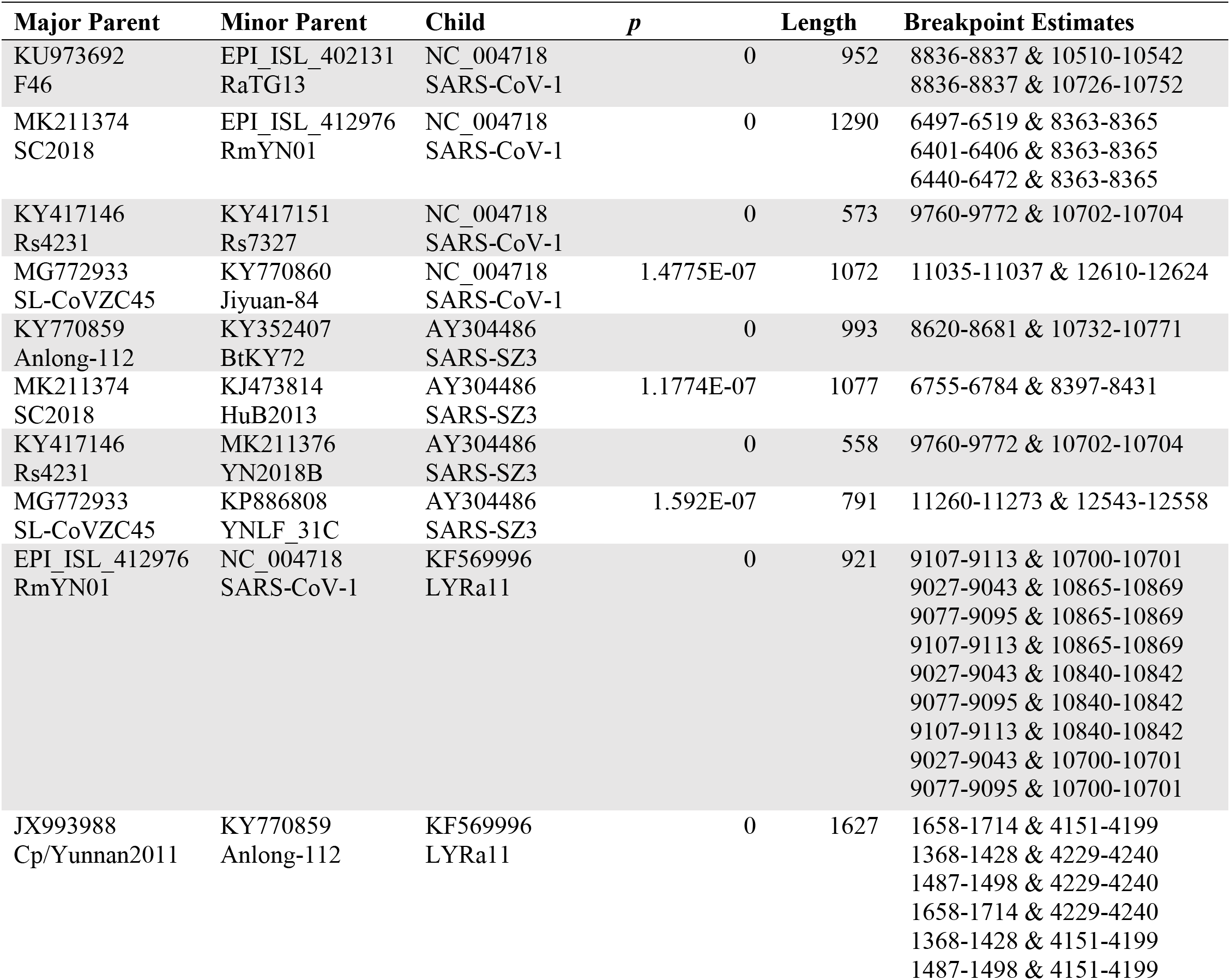
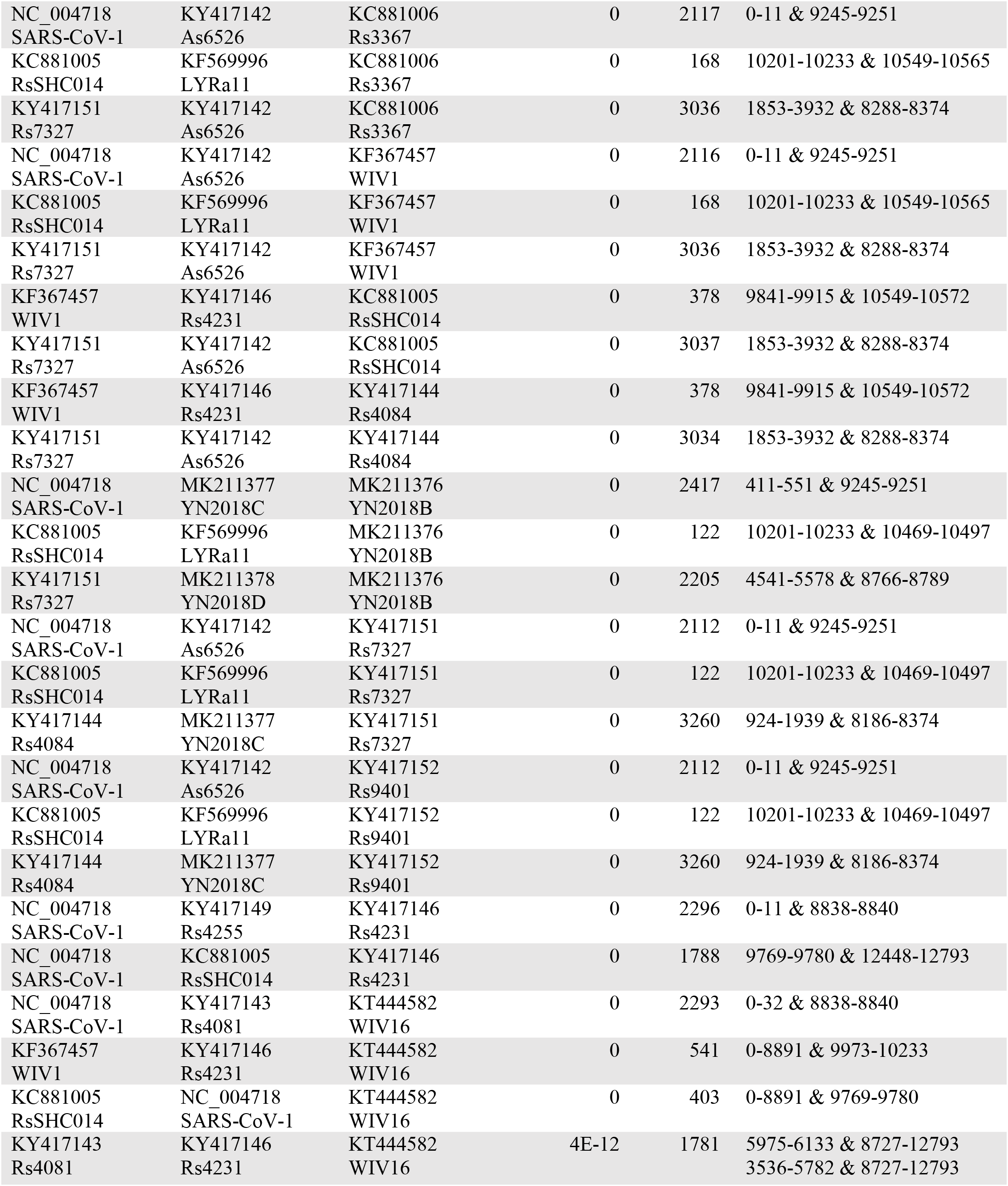
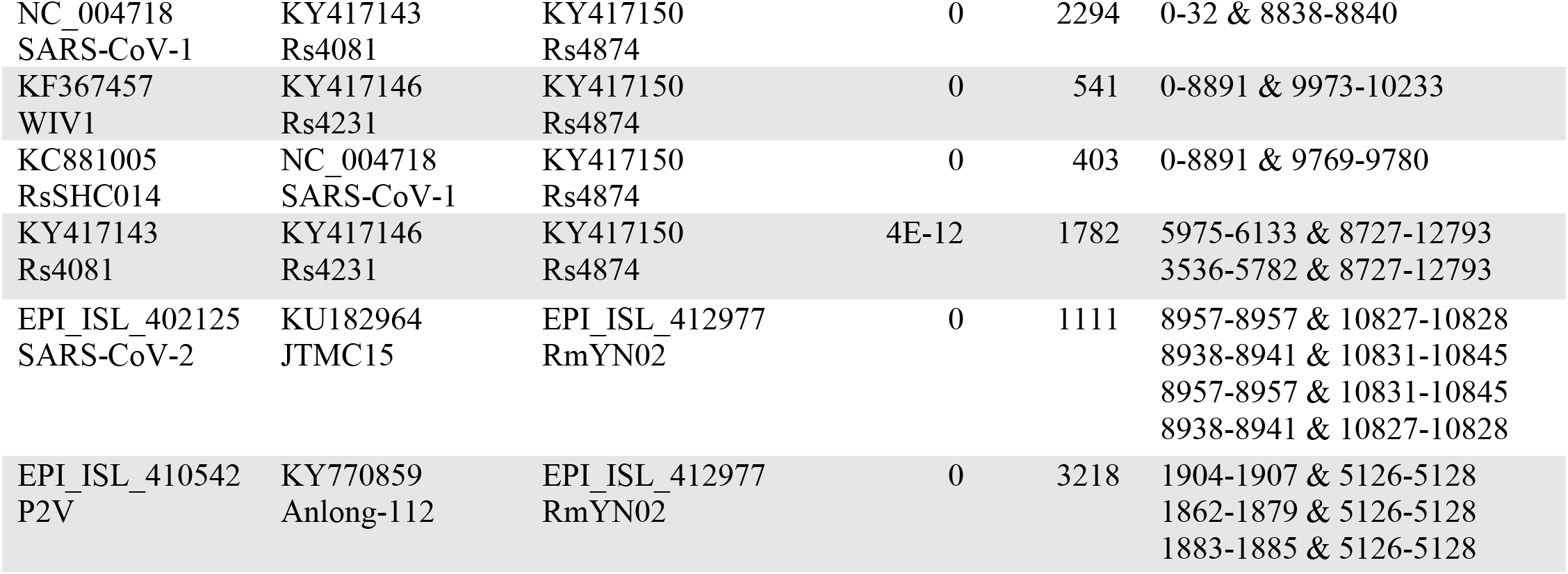
Recombination breakpoints detected in ACE2-using Lineage 1 viruses by the program 3SEQ. Each recombinant Lineage 1 virus was set as the child sequence, and the parental sequences between the breakpoints identified (minor parent) and on either side (major parent) are listed. The *p*-value indicates the level of significance indicated by 3SEQ. Breakpoint estimates are given as ranges, and the minimum length of the recombinant region between these breakpoints is given. Numbering is relative to the alignment, which begins at SARS-CoV-2 nucleotide 12,681. When 3SEQ identified more than one set of breakpoint estimates, all were included in the table. Each recombinant region was further analyzed separately for more breakpoints within, since 3SEQ identifies only one at a time.

Maximum clade credibility trees were built using TreeAnnotator and visualized with FigTree with branches scaled by distance. Posterior probabilities are shown on the preceding branch for each node and probabilities for nodes near the tips of the tree were removed for visual clarity as the exact reconstruction of the most recent divergence events are not within the scope of this study and bear no impact on the interpretation of evolutionary events deeper within the tree.

Finally, for time-calibrated phylogenies, we minimized the effect of recombination on our estimates by using regions of the genome that were free of recombination for the 13 Lineage 1 sequences of interest (further detailed below). In place of RdRp we used Region A, and in place of RBD we used Region E. These regions were determined to be completely breakpoint free for all sequences using 3SEQ. We started by adding tip dates to Region A and used a strict molecular clock with a normally distributed prior informed from estimates derived in Boni et al. (mean 5.5e-4, sd 5.5e-5) [39]. The prior distribution for the coalescent population size was set to lognormal with mean 1 and standard deviation 10 to help with convergence, as the default of 1/X is an improper prior. Our phylogenetics and time estimates are in accordance with those proposed by Boni et al [39]. As the substitution rate in the spike gene is undoubtedly higher than in RdRp, the same clock rate prior could not be used for the Region E time-calibrated phylogeny because the divergence dates would not be comparable. Instead, we assumed the age of the root of this tree should be approximately the same as the age of the Region A tree and fixed the tree height to match the posterior estimate of the tree height for Region A (770 years before present, 1250 AD). This was done by adding a monophyletic MRCA prior to all taxa with a Laplace distribution with mu 1250 and scale 0.1. To account for lineage-specific substitution rates, we also tested a relaxed lognormal clock model.

### Screening for recombination using detection algorithms

We restricted our search for recombination breakpoints to the region of sequence beginning 750 base pairs upstream from RdRp (SARS-CoV-2 nucleotide 12,681) through the end of S2 (through SARS-CoV-2 nucleotide 25,176). There are undoubtedly other breakpoints outside of this region, but since our analysis focuses primarily on RdRp and the spike, the recombination events elsewhere in the genome are outside the scope of this study. We used the program 3SEQ [76] to test the 13 putative recombinants within Lineage 1 (SARS-CoV-1, SARS-SZ3, LYRa11, Rs3367, WIV1, RsSHC014, Rs4084, YN2018B, Rs7327, Rs9401, Rs4231, WIV16, Rs4874) and RmYN02 individually. If breakpoints were found, each subregion on either side of the breakpoint was assessed separately to fine-tune our assessments until no further breakpoints were identified. We did not test any of the remaining sequences for recombination. We were able to identify six regions across all 13 recombinants that appear to be free of recombination and chose these for further phylogenetic analysis (above). The topologies of regions A and E are not significantly different from the topologies of RdRp and the RBD, respectively, suggesting that our use of RdRp and RBD phylogenies in Figures 1, 2, and 5 is a sufficient representation despite some minor evidence of recombination (*e.g*., LYRa11).

### Cell culture and transfection

BHK and 293T cells were obtained from the American Type Culture Collection and maintained in Dulbecco’s modified Eagle’s medium (DMEM; Sigma–Aldrich) supplemented with 10% fetal bovine serum (FBS), penicillin/streptomycin and L-glutamine. BHK cells were seeded and transfected the next day with 100ng of plasmid encoding hACE2 or an empty vector using polyethylenimine (Polysciences). VSV plasmids were generated and transfected onto 293T cells to produce seed particles as previously described [8]. CoV spike pseudotypes were generated as described in [77] and transfected onto 293T cells. After 24h, cells were infected with VSV particles as described in [78], and after 1h of incubating at 37 °C, cells were washed three times and incubated in 2 ml DMEM supplemented with 2% FBS, penicillin/streptomycin and L-glutamine for 48 h. Supernatants were collected and centrifuged at 500*g* for 5 min, then aliquoted and stored at −80 °C.

### Western blots

293T cells transfected with CoV spike pseudotypes (producer cells) were lysed in 1% sodium dodecyl sulfate, 150mM NaCl, 50 mM Tris-HCl and 5 mM EDTA and centrifuged at 14,000*g* for 20 minutes. Pseudotyped particles were concentrated from producer cell supernatants that were overlaid on a 10% OptiPrep cushion in PBS (Sigma–Aldrich) and centrifuged at 20,000*g* for 2h at 4 °C. Lysates and concentrated particles were analyzed for FLAG (Sigma–Aldrich; A8592; 1:10,000), GAPDH (Sigma–Aldrich; G8795; 1:10,000) and/or VSV-M (Kerafast; 23H12; 1:5,000) expression on 10% Bis-Tris PAGE gel (Thermo Fisher Scientific).

### Cell entry assays

Luciferase-based cell entry assays were performed as described in [8]. For each experiment, the relative light unit for spike pseudotypes was normalized to the plate relative light unit average for the no-spike control, and relative entry was calculated as the fold-entry over the negative control. Three replicates were performed for each CoV pseudotype.

### Structural modeling

RBDs were modeled using Modweb [79]. Modeled RBDs were docked to hACE2 by structural superposition to the experimentally determined interaction complex between SARS-CoV-1 RBD and hACE2 (PDB 2ajf) [41] using Chimera [80].

## Acknowledgements

We also thank three anonymous reviewers who provided thoughtful and robust suggestions that substantially improved this manuscript. The research reported in this publication was supported by the National Institute of Allergy and Infectious Diseases of the National Institutes of Health under Award Number R01AI149693 (PI Anthony). ML and VJM are supported by the Intramural Research Program of the National Institute of Allergy and Infectious Diseases (NIAID), National Institutes of Health (NIH). GL and KC are supported by National Institutes of Health (NIH) grant U19AI142777. This study was also made possible by the support of the American people through the United States Agency for International Development (USAID) Emerging Pandemic Threats PREDICT project, GHN-A-OO-09-00010-00 (PI Mazet) and AID-OAA-A-14-00102 (PI Mazet). The content is solely the responsibility of the authors and does not necessarily represent the official views of the U.S. Government.

## Statement of Data Availability

All sequences have been submitted to GenBank and alignments used for phylogenetics are included as supplementary materials.

## Supplementary Materials

**Supplementary Figure S1.**
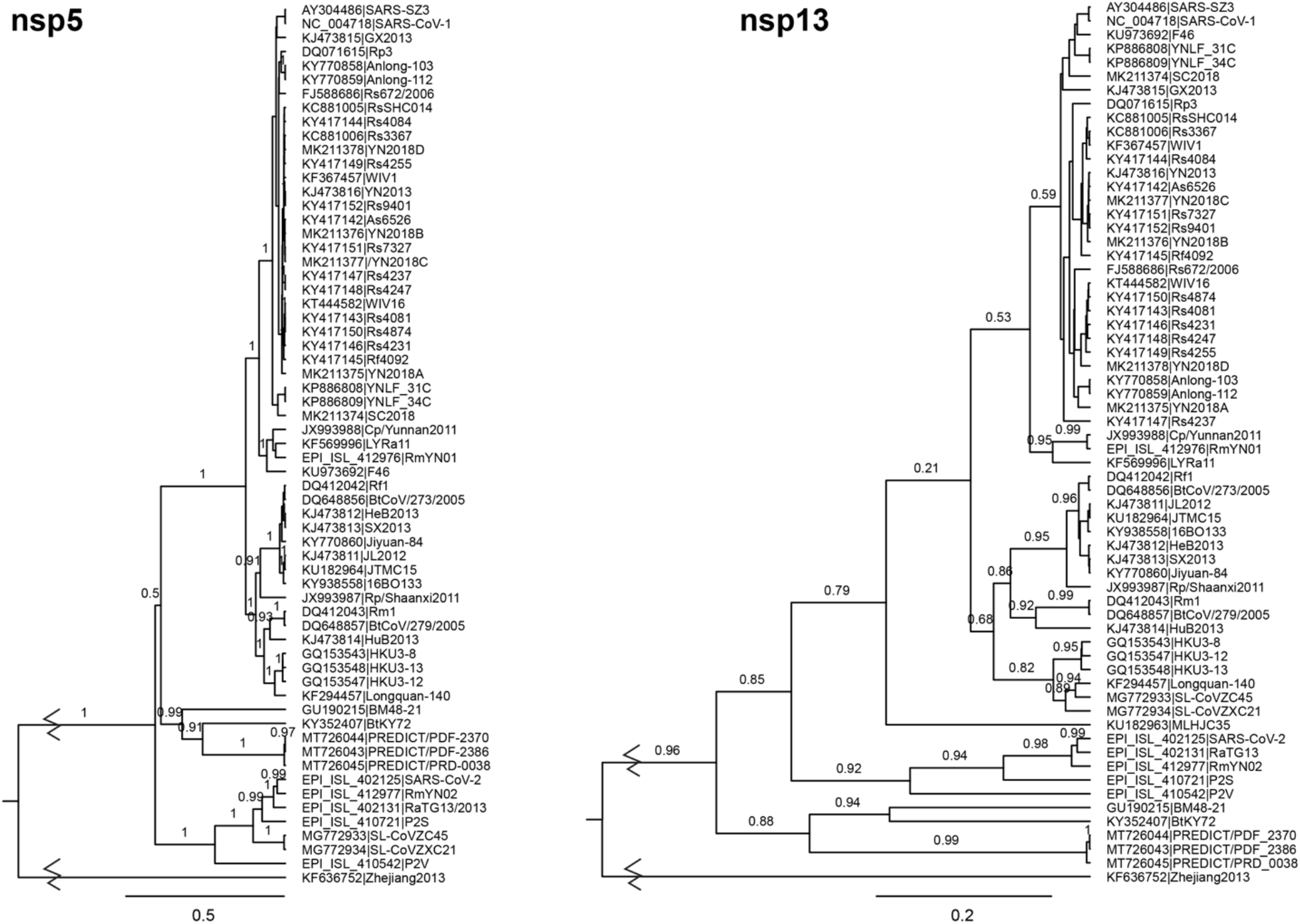
Phylogenetic trees of additional orf1ab genes used for taxonomic classification. To investigate the robustness of the position of Lineage 4 in the RdRp phylogeny, we also constructed phylogenies of nsp5 (3CLpro) and nsp13 (HEL1 core) using identical methods to those used to generate Figure 1. While nsp5 supports the topology we observed for RdRp (nsp12), nsp13 supports the positioning of Lineage 4 at the base of the tree instead. Because of the deep time scale and relatively few sequences used to construct these trees, we must interpret hypotheses that depend on the branching order with caution. The topology is robust to the inclusion or exclusion of the *Hibecovirus* sequence root (data not shown). This pattern of inconsistency was also found for nsp14 and nsp15, with nsp14 matching the topology with Lineage 4 in an intermediate position and nsp15 matching the topology with Lineage 4 at the base (data not shown). The roots of the trees were shortened for clarity.

*Supplementary File 1. Excel spreadsheet of ACE2 amino acid alignment for host species of ACE2-using and non-ACE2-using viruses*. Host ACE2 sequences involved in interfacial interactions with the RBD of SARS-CoV-1 and SARS-CoV-2 are shown for human, civet (*Paguma larvata*), pangolin (*Manis javanica*), and species of bats that are known to both harbor ACE2-binding and non-ACE2-binding viruses (*Rhinolophus sinicus*) or only non-ACE2-binding viruses (*Rhinolophus macrotis, pearsonii, pusillus, ferrumequinum*). The ACE2 sequence from the African bat species from which the PDF-2370 sample was taken is unidentified and also shown. At the time of publication, the ACE2 sequence of *Rhinolophus affinis* was not available. GenBank accession numbers for each sequence are provided.

Distance in angstroms to the nearest SARS-CoV-1 (row 14) or SARS-CoV-2 (row 15) residues are shown and color coded according to the legend in row 18. Residues in hosts of non-ACE2-binders that differ from hosts of ACE2-binders (human, civet, pangolin, and *R. sinicus*) are outlined with black boxes.

*Supplementary File 2. Alignments used for building all phylogenetic trees included in this study*. Alignment files are provided in FASTA format and are named according to the Figure containing the phylogeny constructed from each one.

